# A ratiometric pH sensor for Gram-positive and Gram-negative bacteria

**DOI:** 10.1101/2025.09.09.675137

**Authors:** Dorothea Kossmann, Aya Iizuka, Nina Khanna, Pablo Rivera Fuentes

## Abstract

Fluctuating environments can lead to phenotypic heterogeneity within a monoclonal bacterial population, especially in response to antibiotics or the human immune system. Methods are required to analyze the physiology of single cells to understand how individual cells interact with their environment and adapt to pH stress. We report a ratiometric, fluorescent probe to sense cytoplasmic pH in bacteria. Our probes are based on hemicyanine dyes and are taken up into both Gram-positive and Gram-negative bacteria. The probes react preferentially with OH^-^ over other nucleophiles in biological systems. The response to pH changes is reversible and rapid, allowing for the real-time tracking of pH fluctuations. The sensing of these probes was tuned to allow for monitoring fluctuations around neutrality and biologically relevant acidifications. These probes were validated for cytoplasmic pH sensing in *Escherichia coli*, *Staphylococcus epidermidis*, and a clinically isolated methicillin-resistant *Staphylococcus aureus* (MRSA) strain. Furthermore, the probes enabled the identification of pH-sensitive phenotypes and monitored phagocytosis of virulent clinical strains in immune cells. Our probes are a promising tool for detecting phenotypic heterogeneity within bacterial populations and may help unravel the physiological state of resistant or persistent strains of clinical relevance.

## Introduction

The rise of bacterial antimicrobial resistance is a significant public health threat of the 21st century.^1^ However, it is becoming more evident that even genetically susceptible bacteria may survive antibiotic exposure, and full sterilization is almost never achieved.^2–4^ This antibiotic tolerance differs from resistance in that it does not involve genetic changes, but instead relies on physiological adaptations, such as altered metabolism,^5,6^ growth phase,^7^ and enhanced stress responses.^8^ This phenotypic heterogeneity is modulated by environmental factors, such as heat, acid, antibiotics, and hyperosmotic stress, in addition to stochasticity within biological processes.^9–11^ During infections, the complexity of the host environment induces diverse phenotypic states within genetically identical bacterial populations,^12^ making it difficult to predict individual cell behavior.^13^ Unlike resistance, persistence is difficult to identify due to the absence of reliable markers and the transient nature of underlying physiological states.^4,14,15^ Many phenotype identification methods are time-intensive, such as diffusion assays^16^, colony growth heterogeneity, and bacterial colonies’ lag times, and do not allow for studying phenotypic changes in real-time.^17,18^ In contrast, fluorescent tools enable rapid, non-invasive single-cell studies, revealing phenotypic heterogeneity in complex biological systems.^19^

Bacterial physiology can be studied on several levels, including gene expression, translation, and metabolism,^20^ but one particularly important physiological parameter is intracellular pH.^21^ It affects protein function, bioenergetics, gene expression, stress responses, virulence, and phenotypic switches.^21–25^ Neutralophilic bacteria like *Escherichia coli* (*E. coli*) tightly regulate their cytoplasmic pH between 7.4-7.8,^26–28^ even when exposed to harsh external pH values.^27,28^ The regulatory sensitivity to pH becomes particularly relevant during host-pathogen interactions, as bacteria are often internalized by macrophages into acidic phagosomes.^29^ Acidification aims to degrade bacteria but can unintentionally enhance persistence.^24,30^ Persistence is not solely induced by exposure to low pH. Instead, it has been observed that cells exhibiting a persister phenotype have a more acidic intracellular pH compared to clonal cells that succumb to stress conditions.^31^ This observation suggests that intracellular pH could serve as a potential marker for identifying tolerant cells.

Fluorescent protein sensors, such as the green fluorescent protein (GFP) derivative pHluorin, are widely used to measure bacterial cytoplasmic pH due to their selectivity, brightness, and ability to target specific cellular locations.^31–34^ However, their performance can be affected by buffer composition, bacterial viability, expression levels,^35^ and oxygen availability, which is often essential for chromophore maturation.^36^ Despite extensive refinements, further optimization is often needed for use across diverse bacterial strains.^37–39^ Small-molecule fluorophores offer an alternative, allowing the imaging of non-genetically modified microorganisms, e. g., those isolated from clinical samples.^40^ However, most pH sensors are optimized for mammalian cells, and their uncharged, lipophilic character is poorly suited for bacterial applications since bacteria preferentially take up positively charged, polar, or zwitterionic molecules.^41,42^ Some xanthene-based dyes, such as the cell-permeable acetoxymethyl (AM) esters of fluorescein derivative BCECF **S1** (p*K*_a_ = 7.0, pH-sensing range: 6.5-7.5) or SNARF-1 (p*K*_a_ = 7.5, pH-sensing range: 7.0-8.0), have been applied in bacterial imaging, including studies on host-pathogen interactions.^43,44^ Their pH sensing range, however, is narrow and centered near neutrality. More acid-sensitive derivatives like OregonGreen (p*K*_a_ = 4.7, pH-sensing range: 4.0-6.0)^45^ or SNARF-4 **S2** (p*K*_a_ = 6.4, pH-sensing range: 6.0-7.5)^46^ are sensitive to acidifications, but display a limited pH range, and might require the use of two pH probes to cover a broader dynamic range.^47,48^ Especially SNARF-4F **S2** exhibits reduced bacterial uptake, and is usually used for extracellular pH sensing.^49,50^

To address these limitations, we aimed to develop a bacterial pH sensor capable of sensing pH levels within the physiological range of bacteria, including biologically relevant acidification. Our probe design centered on a coumarin-based dye conjugated to an indoleninium electron acceptor. While a structurally related coumarin-hemicyanine scaffold (CouCy) has previously been reported as a mitochondrial pH sensor, its strong hydrophobic interactions with membranes likely prevent bacterial uptake.^51,52^ Similarly, an aggregation-induced ratiometric pH sensor with a large dynamic range has been reported,^24,53^ but its high molecular weight and hydrophobic character limit its application in bacterial imaging. Even though a benzooxazine hemicyanine was reported for bacterial imaging, its pH sensing ability is limited to highly acidic (pH 2-4) environments.^54^ Therefore, while a broad array of pH sensors exists for mammalian cell imaging, this repertoire still lacks a ratiometric sensor specifically designed to meet the unique demands of bacterial imaging.

We aimed to fine-tune the pH sensitivity of CouCy scaffolds to achieve a suitable pH-sensing range for visualizing bacterial heterogeneity. Our redesigned probes exhibit properties that favor accumulation in *E. coli* according to the eNTRy guidelines^51^ and allow for live-cell pH sensing experiments. Furthermore, the dynamic range was tuned to cover pH changes around neutrality as well as biologically relevant acidification. The applicability of our probes was validated in both Gram-positive and Gram-negative bacteria and applied to study pH-sensitive *E. coli* knockout strains and host-pathogen interactions of resistant and persistent patient-derived strains.

## Results and Discussion

### CouCy probes are selective, sensitive, and fast pH sensors

The CouCy probes **1a-c** were synthesized in a condensation reaction of Fischer’s bases **2a-c** and coumarin aldehyde **3** (Figure 1A). We titrated the CouCy probes to assess their pH sensitivity (Figure 1B, Figure S1) and calculated the p*K*_a_ values based on the Henderson-Hasselbalch equation (Table S1). We found that the pH sensitivity can be tuned by the substituent on the indoleninium core. The unsubstituted derivative CouCyH **1a** has a p*K*a value of 9.4, whereas the introduction of electron-withdrawing groups (EWG) like the trifluoromethyl (CF_3_) and nitrile (CN) in CouCyCF_3_ **1b** and CouCyCN **1c** lowered the p*K*_a_ to 7.0 and 6.8, respectively. Since CouCyCN **1b** and CouCyCF_3_ **1c** exhibit a p*K*_a_ value close to the internal pH range of neutrophilic bacteria, we focused on the further characterization of these probes

**Figure 1:**
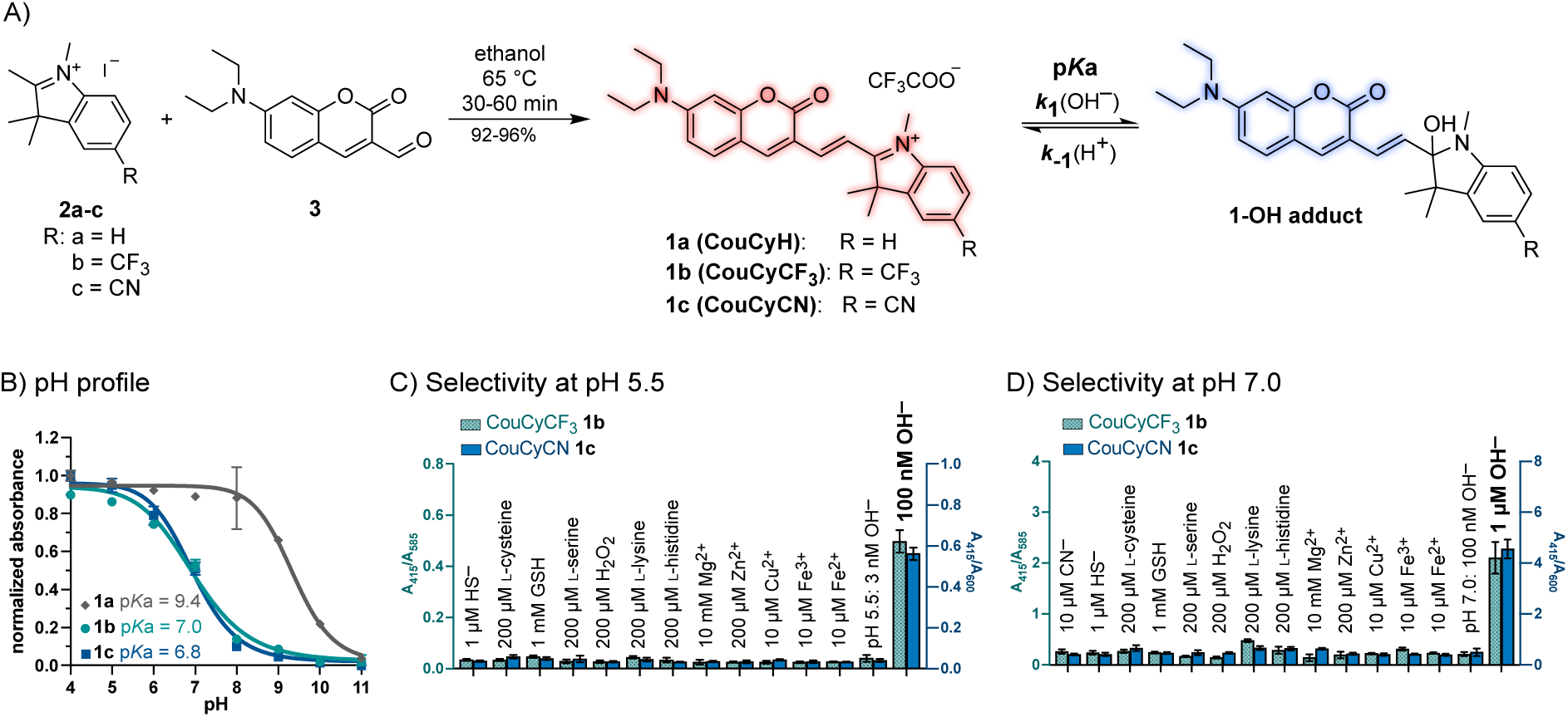
Synthesis and characterization of CouCy’s. A) Condensation reaction of indoleninium **2a-c** and coumarin aldehyde **3** to yield the CouCy derivative **1a, 1b,** and **1c** and equilibrium of the pH sensing reaction. B) pH profile of CouCyH **1a** (p*K*a = 9.4), CouCyCF3 **1b** (p*K*a = 7.0), and CouCyCN **1c** (p*K*a = 6.8). The probe (5 µM) was incubated for 60 min at 37 °C with defined pH values. As pH buffer citric acid and Na2HPO4 (pH 2–8), NaHCO3 and Na2CO3 (pH 9–11), or NaOH and KCl (pH 12–13) were used. Absorbance was normalized to the absorbance maximum *A*max (**1a**: 570 nm; **1b**: 584 nm; **1c**: 600 nm) and interpolated as sigmoidal curves. The p*K*a values were calculated with the Henderson-Hasselbalch equation 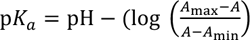, with *A*max as red absorbance maxima and *A*min as blue absorbance maxima (**1a**: *A*440 and *A*568; **1b**: *A*584 and *A*415; **1c**: *A*600 and *A*415). C,D) Selectivity assays of CouCyCF3 **1b** (green bars; *A*415/*A*585) and CouCyCN **1c** (blue bars; *A*415/*A*600) at pH 5.5 (C) and pH 7.0 (D). Absorbance spectra were measured in the presence of 1 µM NaSH, 1 mM GSH, 200 µM H2O2, 200 µM L-cysteine, 200 µM L-serine, 200 µM L-lysine, 200 µM L-histidine, 10 µM CN^-^ (only at pH 7.0), 10 mM Mg(NO3)2, 200 µM Zn(NO3)2, Cu(NO3)2, Fe(NO3)3 and Fe(SO4)2. Probe (10 µM) was dissolved in 10x PBS (pH 5.5 or 7.0) and incubated with the analytes for 30 min at 37 °C and 180 rpm. All data are mean values of triplicates and error bars represent the standard deviation (SD).

Based on structurally similar molecules, we assumed the CouCy scaffold to undergo a nucleophilic attack by hydroxide ions (OH^-^), resulting in a shorter conjugation system as illustrated in Figure 1A.^52^ To investigate the pH sensing mechanism, we measured ^1^H NMR spectra of CouCyCN **1c** in deuterated acetonitrile (CD_3_CN) with NaOD. In the absence of NaOD, a single signal for the two methyl groups (C13) on the indoleninium was observed (Figure S2). After incubation with NaOD, this signal splits into diastereotopic singlets (13’ and 13’’), indicating the formation of a stereocenter nearby, with the methyl group (C9) on the indoleninium nitrogen displaying an upfield shift. In addition, the methine groups on C7 and C8 show a *trans*-coupling constant before (*^trans^*^,3^*J*_7,8_ = 15.5 Hz) and after NaOD treatment (*^trans^*^,3^*J*_7,8_ = 16.0 Hz). Thus, an attack by OH^-^ on the electrophilic Michael acceptor can be excluded. This regioselectivity differs from that of previously reported phosphine nucleophiles, which typically attack CouCy scaffolds at the bridging double bond (C7).^55^ The difference in regioselectivity of the nucleophilic attack might be explained by the Hard Soft Acid Bases (HSAB) principle. Whereas OH^-^ ions are hard nucleophiles, phosphines (PR_3_) are softer and react with the soft, electrophilic position on C7.^56^ Overall, the NMR data suggest an OH^-^ attack on the electrophilic *sp*^2^ carbon on the indoleninium, consistent with the literature on a similar scaffold.^52^ Notably, the EWG group (R = CN or CF_3_) on the indoleninium influences both the sensitivity to OH^-^ and the regioselectivity of the nucleophilic attack. At high OH^-^ concentrations (pH ≥ 10), the lactone of the coumarin core can be hydrolyzed, resulting in a longer conjugation pathway and red-shifted absorbance (Figure S1).^57^ Such a reactivity was observed with CouCyH **1a**, indicated by a red shift in the absorbance maxima *A*_max_ (pH<10: *A*_max_ = 570 nm; pH>10: *A*_max_ = 650 nm; Figure S1). Thus, the OH^-^ regioselectivity depends on the trigonal carbon’s electrophilicity, with more electron-rich CouCy indoleninium prone to undergo a competitive nucleophilic attack on the coumarin lactone. This observation further supports the importance of the electronic tuning introduced with the CN and CF_3_ groups.

We investigated the sensing kinetics of CouCyCF_3_ **1b** and CouCyCN **1c** *in vitro* under pseudo-first-order conditions with an excess of OH^-^ or H^+^, respectively (Figure S3). We found that the OH^-^ attack on CouCyCN **1c**, with an observed pseudo first-order rate constant *k*_1,obs_ = 0.15 s^-1^, is twice as fast as on CouCyCF_3_ **1b**, with a *k*_1,obs_ = 0.09 s^-1^. This observation aligns with the reactivity trend based on EWG strength. These experiments demonstrate that the sensing reaction is reversible and complete within 1-2 min, allowing for dynamic monitoring of intracellular pH fluctuations with good time resolution.

To assess the selectivity for OH^-^, we analyzed the ratiometric absorbance change of CouCyCF_3_ **1b** and CouCyCN **1c** in the presence of other nucleophiles at pH 5.5 and 7.0 (Figure 1). We used the ratio *A*_blue_/*A*_red,_ where the blue signal corresponds to the conjugate base CouCy-OH and the red signal to the parent CouCy (conjugate acid). Since the base is formed upon nucleophilic attack, *A*_blue_ in the numerator emphasizes the extent of CouCy-OH formation, and a change in *A*_blue_/*A*_red_ ratio assesses the reactivity with other nucleophiles. The following analytes were used in biologically relevant concentrations or excess: 1 µM HS^-^, 1 mM reduced glutathione (GSH), 200 µM H_2_O_2_, 200 µM of the nucleophilic amino acids L-cysteine, L-serine, L-lysine, L-histidine, 10 mM Mg^2+^, 10 µM Cu^2+^, 200 µM Zn^2+^, 10 µM Fe^2+^ and Fe^3+^ and 10 µM CN^-^ (see SI for comment on concentration ranges). The selectivity assays revealed that CouCyCF_3_ **1b** and CouCyCN **1c** are selective to OH^-^ ions over other nucleophiles or ions of biological relevance (Figure 1C,D). Based on reported indoleninium-based sensor probes, CN^-58,59^, HS^-60,61^, and GSH^62,63^ are the most competitive nucleophiles to OH^-^. Whereas cyanogenic bacteria like *Pseudomonas aeruginosa* can produce CN^-^ in the µM range, CN^-^ is commonly absent in most bacteria due to its toxicity.^64^ Thus, we assume no interference by CN^-^ ions on our pH sensing in *E. coli*, *Staphylococcus* (*S.) epidermidis*, or *S. aureus*. The reported H_2_S concentration in bacteria is the low micromolar range (*E. coli:* 2.5 µM; *S. aureus*: up to 38 µM).^65^ In an OH^-^/HS^-^ competition study, we found that CouCyCF_3_ **1b** is more selective than CouCyCN **1c**. The pH sensing ability of CouCyCF_3_ **1b** remained unaffected in the presence of 10 µM HS^-^ (Figure S4), whereas CouCyCN **1c** undergoes competitive nucleophilic attack of HS^-^ indicated by a change in the *A*_blue_/*A*_red_ ratio. At higher HS^-^ concentrations (≥ 100 µM), both probes exhibit competitive reactivity, particularly above pH 7.0, where the fraction of deprotonated HS^-^ increases. Given that physiological concentrations of H_2_S are unlikely to surpass 100 µM and our probes are largely intended to be used for sensing acidification, the interference of HS^-^ in intracellular pH sensing is negligible. In a second thiol-pH competition study, we assessed the reactivity of CouCy dyes with GSH. In most Gram-negative bacteria, GSH is present in the mM range, but is absent in Gram-positive species.^65^ We found that below pH 8.0 and at concentrations from 0.01 to 5 mM, GSH had a minimal impact on the ratiometric signals of CouCyCF_3_ **1b** (Figure S4). In contrast, CouCyCN **1c** showed an increased susceptibility to nucleophilic attack by GSH, as previously described for HS^-^. These measurements revealed that CouCyCF_3_ **1b** is more selective for pH sensing and indicate under which conditions the effect of HS^-^ and GSH fluctuations could be safely ignored, which is mostly at acidic pH (pH < 7.00).

### CouCy sensors reliably detect pH changes in live Gram-negative and Gram-positive bacteria

To evaluate the pH-sensing capability of CouCyCF_3_ **1b** and CouCyCN **1c** in live cells, we performed intracellular pH calibration experiments using *E. coli* and *S. epidermidis*. The bioavailability of small-molecule probes in bacteria depends on uptake, distribution, and efflux pathways.^41^ Especially, staining of Gram-negative bacteria is challenging due to the permeability barrier of the outer membrane.^66^ However, the CouCy scaffold has physicochemical properties that favor accumulation in *E. coli*, like a small size (<600 Da), a positive charge, low globularity (≤0.25), and rigidity due to the conjugation system (Table S2).^51^ Furthermore, the dual-wavelength response of CouCy probes enables ratiometric fluorescence measurements, providing a readout that is independent of probe concentration and instrument fluctuations, thereby increasing accuracy and sensitivity.^67^ Such self-calibration is particularly advantageous for bacterial imaging, where the bioavailability of small-molecule probes is modulated by uptake and efflux and may fluctuate depending on the bacteria’s environment and physiological state.^41^

We used *I*_red_/*I*_blue_ for ratiometric analysis, as it provided a 10 times broader dynamic range compared to the inverse *I*_blue_/*I*_red_ ratio and exhibits better sensing ability in the desired acidic range with an improved signal-to-noise ratio, as the signal of the conjugate acid (parent CouCy) dominates the ratio (Figure S5).

To evaluate the uptake efficiency, we incubated the Gram-negative *E. coli* K12 with 5 µM, 2 µM, or 1 µM CouCyCF_3_ **1b** and found strong intracellular staining at all concentrations (Figure S6), suggesting efficient intracellular accumulation. Notably, bacterial staining was observed immediately, without the need for extended incubation time. The ratiometric signal remained constant across concentrations, demonstrating the self-calibration and concentration independence of our probes (Figure S6).

Despite the poor emission quantum yield ø_em_ (**1b:** 0.8%^55^; **1c**: 1.4%), extinction coefficients ε (**1b**: 63 × 10^3^ M^-1^cm^-1^; **1c**: 51 × 10^3^ M^-1^cm^-1^) and low brightness ε × ø_em_ (**1b**: 509 M^-1^cm^-1^; **1c**: 715 M^-1^cm^-1^) of CouCy dyes in aqueous solution, we observed good intracellular signal, suggesting some degree of fluorogenicity. We propose that this effect might be related to the polarity-dependent photophysical properties of CouCy dyes, which exhibit low quantum yields in aqueous media but increased emission in apolar intracellular environments (Figure S7, Table S3). Both probes exhibit an increase in blue emission and a decrease in red emission intensity as the polarity decreases. Interestingly, in very apolar media, which cover the polarity range of bacteria (dielectric constant ε < 10),^68–70^ CouCyCN **1c** exhibits intense blue and almost no red emission intensities, whereas CouCyCF_3_ **1b** shows intense blue and a moderate red emission intensity. This finding suggests that CouCyCF_3_ **1b** exhibits photophysical properties that are advantageous for bacterial imaging. This effect, albeit useful for imaging, could pose a challenge for pH quantification if the brightnesses of the hydroxylated and parent CouCy dyes varied strongly with polarity. Fortunately, we found that this effect is negligible within the polarity range of bacteria, as the ratiometric readout *I*_red_/*I*_blue_ remains constant in highly apolar environments (Figure S7). Due to the polarity dependence, the dynamic range in aqueous media (Figure S5) may not accurately represent the pH sensing ability of the probes in cells. Thus, we calibrated these sensors directly in live cells using carbonyl cyanide *m*-chlorophenyl hydrazone (CCCP; 250 µM), a chemical inhibitor of oxidative phosphorylation and a mediator of H^+^ influx that uncouples the cellular proton gradient.^71^ CCCP allows to equilibrate the extracellular and intracellular pH, bypassing the ability of bacteria to tightly regulate their intracellular pH.^27,28^

Live-cell fluorescence microscopy of *E. coli* K12 stained with CouCyCF_3_ **1b** (Figure 2) showed a pH-dependent emission shift: under acidic conditions, fluorescence was mainly observed in the red channel (*α*_ex_ = 640 nm), whereas increasing pH led to a shift towards blue emission (*α*_ex_ = 445 nm). Ratiometric images at pH 5.0, 6.0, and 7.0 (Figure 2B) demonstrate clear pH-dependent changes at the single-cell level. Quantitative analysis of the *I*_640_/*I*_445_ ratio revealed a linear response between pH 5.0 and 7.5 (Figure 2C,D), based on two independent biological replicates. Furthermore, we used *S. epidermidis* as a model organism to evaluate the pH sensing of CouCyCF_3_ **1b** in Gram-positive strains. Confocal images showed similar pH-induced emission shifts (Figure S8), and the ratiometric images confirm a linear *I*_640_/*I*_445_ response from pH 5.0 to 7.5 (Figure 2E-G), consistent with the results in *E. coli*. These results demonstrate that CouCyCF_3_ **1b** can be employed to sense pH in both Gram-negative and Gram-positive bacteria.

**Figure 2:**
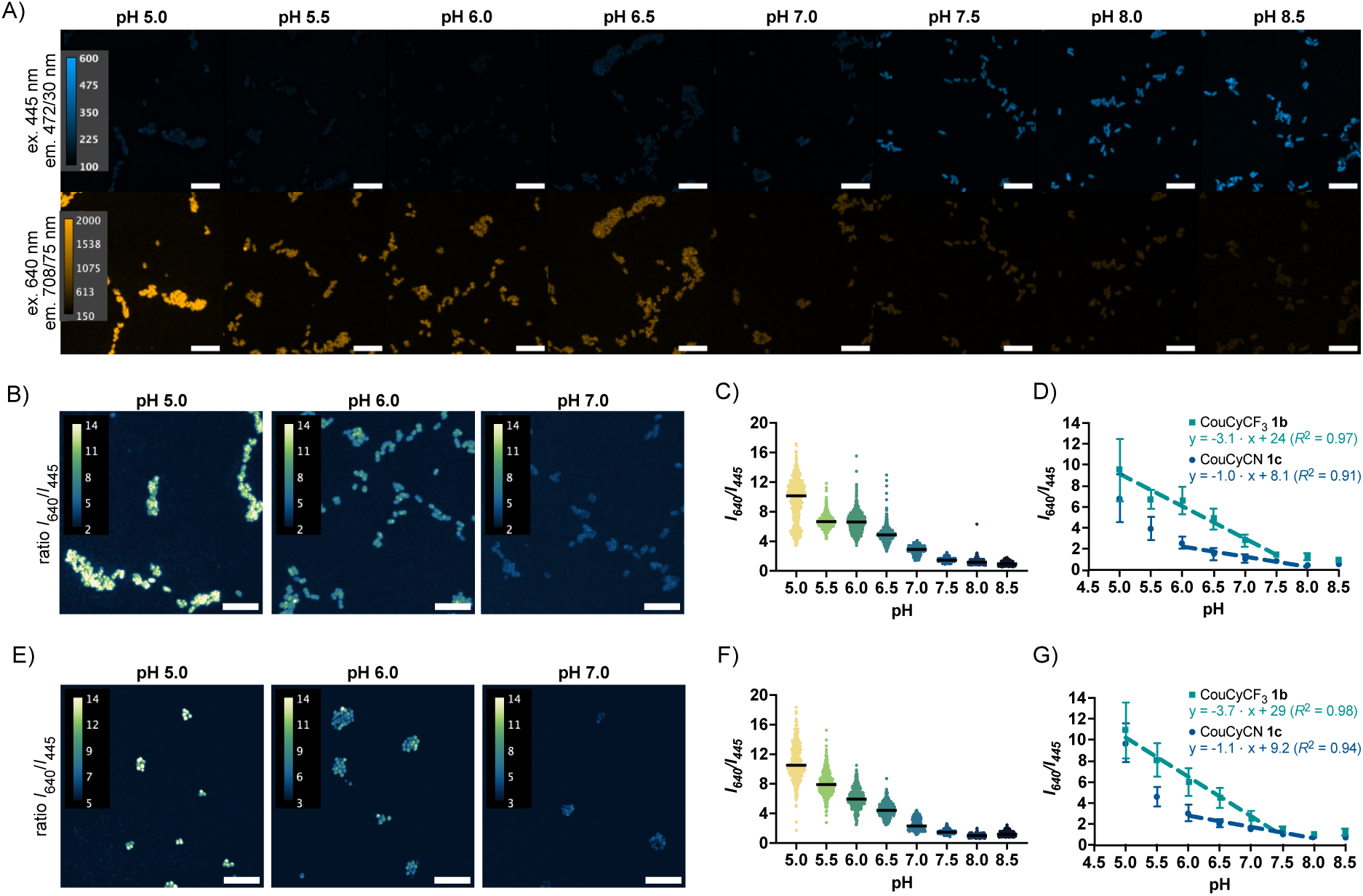
Microscopy images and live-cell pH calibration experiments. A) Confocal microscopy images of *E. coli* treated with CouCyCF3 **1b** (1 µM) for 20 min at 37 °C, followed by a CCCP (250 µM) treatment in PBS (10x, varying pH) for 60 min at 37 °C. B) Representative ratiometric images of *E. coli* treated with CouCyCF3 **1b** (1 µM) and CCCP (250 µM) at pH 5.0, 6.0, and 7.0. C) Scatter plot of *I*640/*I*445 ratio from *E. coli* stained with CouCyCF3 **1b** (1 µM), with *N* = 499, 683, 771, 747, 545, 762, 734, 545 (from left to right), independent single cells or cell clusters from two separate imaging sessions. D) Linear regression of the *I*640/*I*445 mean with SD in *E. coli* K12 stained with CouCyCF3 **1b** (1 µM) or CouCyCN **1c** (1 µM). E) Representative ratiometric images of *S. epidermidis* treated with CouCyCF3 **1b** (1 µM) and CCCP (250 µM) at pH 5.0, 6.0 and 7.0. F) Scatter plot of *I*640/*I*445 ratio with *N* = 238, 352, 388, 339, 234, 310, 189, 220, independent single cells or cell clusters from two separate imaging sessions. G) Linear regression of the *I*640/*I*445 mean with SD in *S. epidermidis* stained with CouCyCF3 **1b** (1 µM) or CouCyCN **1c** (1 µM). Imaging was performed with the following laser setup: ex. 445 nm, em. 472/30 nm, 1.8 mW, 400 ms; ex. 640 nm, em. 708/75 nm, 3.8 mW, 400 ms. Scale bar, 10 µm.

CouCyCN **1c** was also evaluated in both *E. coli* K12 and *S. epidermidis*. Ratiometric fluorescence analysis revealed linear pH-sensing between pH 6.0 and 8.0 in both microorganisms (Figure 2, Figure S9). Notably, the calibration slopes differ significantly, with CouCyCF_3_ **1b** showing steeper slopes (*m* = −3.1 and −3.7) compared to CouCyCN **1c** (*m* = −1.0 or −1.1), indicating higher sensitivity of CouCyCF_3_ **1b** to pH changes. Furthermore, the linear pH-sensing range of CouCyCF_3_ **1b** extended to pH 5.0. This difference in dynamic range was further highlighted in pH sensing experiments using flow cytometry. *E. coli* K12 were incubated with either probe (1 µM) in the presence of CCCP (250 µM) across pH 5.0 to 8.0. The bacterial populations stained with CouCyCF_3_ **1b**, showed a clear pH-dependent separation, whereas samples stained with CouCyCN **1c** showed less distinct separation across the same pH range (Figure S10). The enhanced dynamic range and increased sensitivity of CouCyCF_3_ **1b** suggest its suitability for high-throughput applications such as fluorescence-activated cell sorting (FACS) to visualize single-cell phenotypes in the analysis of multiple cells. The superior performance of CouCyCF_3_ **1b** may be attributed to its photophysical properties, particularly the polarity-dependent emission intensities, which exhibit strong emission signals at both emission maxima within the polarity range of bacterial cells (Figure S7).

To assess the performance of our probes relative to established pH sensors, we conducted experiments using reported ratiometric pH sensors with comparable p*K*_a_ values. We chose the ratiometric biosensor mCherry-pHluorin, which combines the pH-insensitive mCherry with the pH-sensitive GFP derivative pHluorin. We utilized the plasmid pSCM001 (Addgene: 124605),^34^ which encodes the mCherry-pHluorin fusion protein under an arabinose-inducible promoter. Confocal microscopy images (Figure S11) show the expected pH-induced trend, characterized by increasing pHluorin emission (*α*_ex_ = 488 nm) while mCherry emission (*α*_ex_ = 561 nm) remained constant.^34^ Ratiometric analysis of *I*_488_/*I*_561_ revealed a linear pH sensing trend between pH 6.5-8.0 as reported in the literature (Figure S11).^31,34^ Notably, CouCyCF_3_ **1b** showed a larger dynamic range (pH 5.0-7.5), better sensing ability at low pH, and higher sensitivity indicated by a larger slope (*m* = 3.1) compared to mCherry-pHluorin (*m* = 2.2).

In addition to the protein sensor, we conducted experiments with the commercial small-molecule probe BCECF-AM **S1** with a p*K*_a_ of 6.9 (Figure S12).^72^ This compound has been successfully applied in bacterial imaging, including studies on host-pathogen interactions.^43,44^ We stained *E. coli* K12 with BCECF-AM **S1** (1 µM) and found a linear *I*_488_/*I*_445_ ratio change between pH 6.5 and 8.5 (Figure S12). The fluorescein derivative showed a high intracellular emission but high background signals, likely resulting from spontaneous hydrolysis in aqueous solutions.^73^ Furthermore, BCECF **S1** displays a narrower sensing range than CouCyCF_3_ **1b**. The pH-sensitive emission change of xanthene-based dyes is based on the protonation of the xanthene core. The dianion, as conjugate base, is the main fluorescent species, whereas the protonated, conjugate acid species exhibits a weaker donor-acceptor system with diminished fluorescence.^74^ In contrast, our CouCyCF_3_ **1b** sensor exhibits strong fluorescence from both its conjugate acid and base, as the molecule functions as two interconverting fluorophores, each dominating at different pH values. This feature enables ratiometric sensing with higher signal intensity across a larger range.

In addition, we aimed to conduct pH sensing experiments with the commercial small molecule SNARF-4F-AM **S2** (p*K*_a_ = 6.4), which exhibits a more acidic pH sensing range (pH 6.0-7.5).^46^ However, the low emission intensity observed indicated insufficient uptake of the probe (Figure S13). This finding is not surprising, as SNARF-4F **S2** is primarily used for measuring extracellular pH in bacterial studies.^49,50^ In conclusion, in comparison to commonly used small-molecule pH sensors, our probes exhibit good bacterial accumulation, a broader dynamic range, and increased sensitivity to detect pH changes.

### CouCyCF_3_ detects decreased acid tolerance in bacteria lacking cyclopropane fatty acids

Cyclopropane fatty acids (CFAs) are linked to stress protection and acid tolerance of bacteria.^75–77^ These fatty acids are synthesized from monounsaturated fatty acids catalyzed by the CFA synthase (cfaS) as a post-synthetic modification of the phospholipid bilayer (Figure 3A). Elevated CFA levels have been associated with increased membrane fluidity and enhanced acid tolerance in *E. coli* strains.^75,76,78^ Conversely, bacteria that lack CFA in their membranes exhibit increased proton permeability.^77^ Previous studies have assessed the acid sensitivity of CFA synthase knockout (Δ*cfaS*) strains using microelectrode-based H^+^ flux measurements^77^ or survival studies based on colony formation on agar plates.^75,76^ We investigated whether our pH probe could visualize the different phenotypes of Δ*cfaS* knockout and parental *E. coli* BW25113 under stress conditions. The acid shock was performed by incubating the cells stained with CouCyCF_3_ **1b** (1 µM) in lysogeny broth (LB) at different pH values (7, 5, 4, and 3).^75,76^ The change of cytoplasmic pH was monitored over time by flow cytometry, allowing for gating acidified cells based on the red (*I*_639_ > 3.5 x 10^3^) and blue (*I*_405_ nm > 1 x 10^3^) emission channels (Figure S14). Consistent with literature reports,^27,28^ both strains maintained a stable intracellular pH under mildly acidic conditions (pH 5 or 4), as less than 1% of the cells were acidified (Figure S14). However, exposure to pH 3 resulted in acidification of the cytoplasm, consistent with lethal acid shock conditions (Figure 3B).^75,76^ Importantly, the Δ*cfaS E. coli* population undergoes a more significant acidification compared to the parental strain, as indicated by the appearance of a more dominant population with an increased *I*_639_/*I*_405_ emission ratio (Figure 3B,C). These results confirm the increased sensitivity of Δ*cfaS* strains to acid stress and demonstrate that CouCyCF_3_ **1b** enables real-time visualization of pH-sensitive phenotypes at the single-cell level.

**Figure 3:**
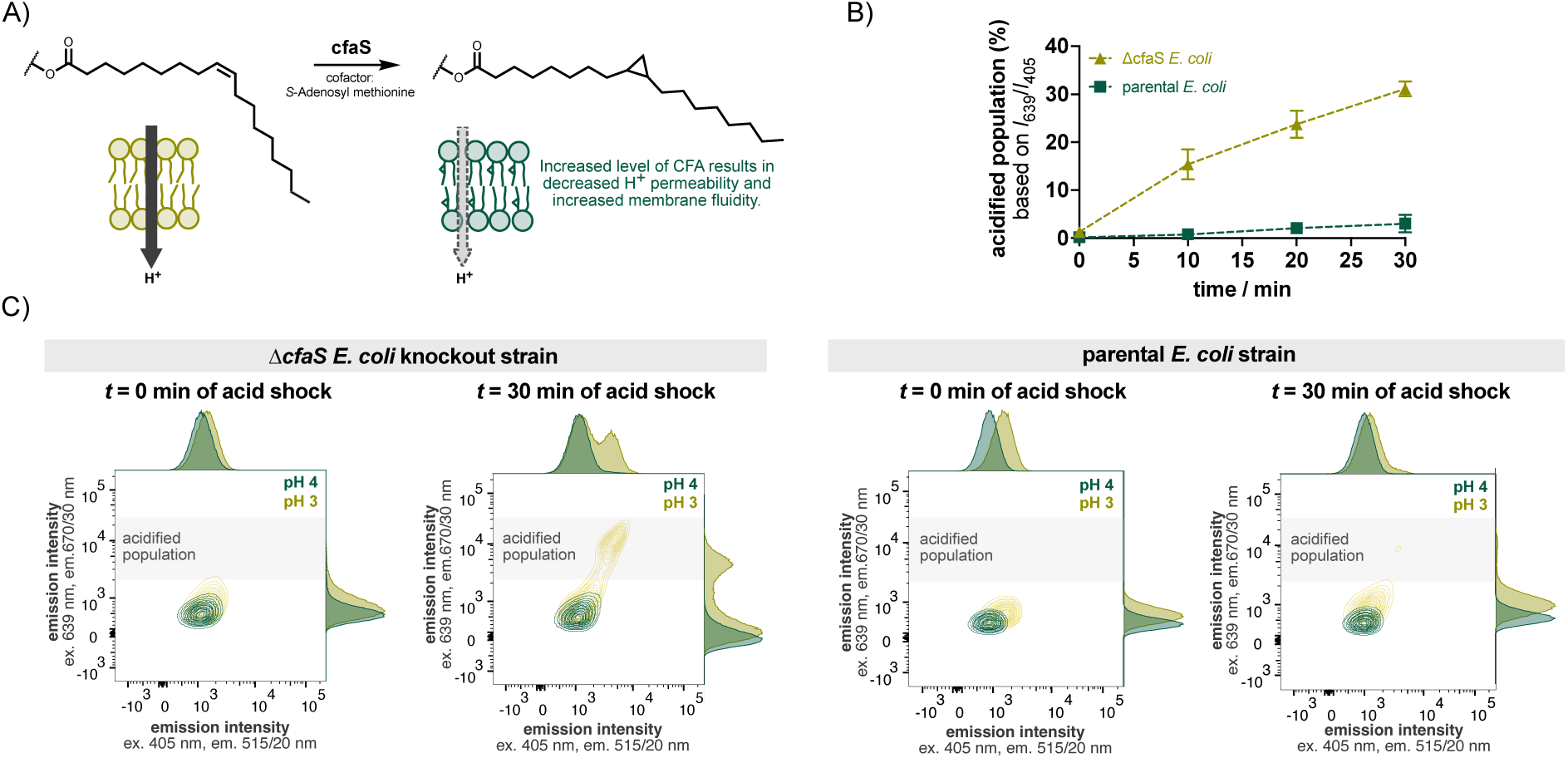
Comparison of pH-sensitive Δc*faS E. coli* knockout strain with the parental *E. coli* BW25113 (Keio Knockout Collection)^79,80^. A) Reaction scheme of the cfaS-catalyzed synthesis of CFA using monounsaturated fatty acids and *S*-adenosyl methionine as cofactor. CFA leads to a lipid bilayer with decreased packing density and increased membrane fluidity, resulting in lower proton membrane permeability.^77,81^ B) Quantitative analysis of Δc*faS* (beige) and parental *E. coli* (green) population under acid shock at pH 3 over 30 min stained with CouCyCF3 **1b**. Acidified cells were gated based on emission intensity of the red (*I*639 > 3.5 x 10^3^) and blue (*I*405 nm > 1 x 10^3^) channels. Data points are means with SD from two biological replicates. C) Flow cytometry analysis representing bacterial populations as contour plots and adjunct histograms at time points 0 and 30 min of the acid shock at pH 3 and 4. Each sample was gated for single bacterial cells.

### CouCy probes can track bacterial acidification in host-pathogen interaction studies

To answer clinically relevant questions, phenotypes should be studied under in-patient-like conditions, as standard laboratory settings may not accurately replicate virulent phenotypes.^12,20,82^ In this context, we evaluated the applicability of the CouCy probes for host-pathogen interaction studies using the laboratory strain *S. epidermidis* and clinical isolates of *S. aureus*. Specifically, a methicillin-resistant *S. aureus* (MRSA, t619915) strain isolated from a prosthetic joint infection, and a methicillin-sensitive *S. aureus* (MSSA, P70) strain from a deep-seated infection and bacteremia were collected from patients at the University Hospital Basel, Switzerland. The clinical *S. aureus* was efficiently stained with CouCyCF_3_ **1b** and CouCyCN **1c**, allowing for the visualization and quantification of intracellular pH using fluorescence microscopy or flow cytometry (Figure S15). Additionally, we assessed the impact of CouCy dyes on bacterial viability. At the working concentration of 1 µM, no cytotoxic effects were observed, as bacterial growth remained unaffected over 24 hours. Higher concentrations (5 µM) induced only a minor reduction in bacterial growth rate (Figure S16), indicating that CouCy dyes are well tolerated under the experimental conditions used.

For infection studies, the laboratory strain *S. epidermidis* or clinical isolates of *S. aureus* were pre-stained with CouCyCF_3_ **1b** (1 µM) before coculturing with THP-1 monocytes in serum-containing medium. A multiplicity of infection (MOI) of 3 was induced to achieve three times more bacterial cells than immune cells. Intracellular bacteria were detected 10 minutes post-infection using confocal microscopy (Figure S17) or imaging-based flow cytometry (*ImageStream*; Figure S18). The increased *I*_red_/*I*_blue_ emission ratio indicates that intracellular bacteria are experiencing acid stress in phagolysosomes, whereas extracellular bacteria maintain their intracellular pH neutral, with a lower *I*_red_/*I*_blue_ ratio. This experiment demonstrates that CouCyCF_3_ **1b** enables the detection of extracellular and intracellular bacteria and can be used to study phagocytosis in immune cells. A widely used fluorescent dye for monitoring phagocytosis is pHrodo™ Red^83^, which exhibits increased emission at low pH and has a broad dynamic range (pH 4-8). Despite its utility, pHrodo™ Red provides only a single-channel readout, limiting its application to qualitative assessments.^84^ In contrast, the ratiometric properties of CouCyCF_3_ **1b** enable quantitative pH measurements, facilitating comparative studies. Thus, we conducted a quantitative intracellular pH comparison study as part of our host-pathogen interaction experiment. To mimic in-patient-like conditions accurately, blood cells from freshly heparinized human whole blood were extracted and cocultured with clinical *S. aureus* strains. The obtained blood cell samples represent a biologically complex mixture, including monocytes, neutrophils, and lymphocytes. As previously demonstrated for the THP-1 cell line, we were able to identify extracellular and intracellular bacteria 10 minutes post-infection (Figure 4A, Figure S19). With flow cytometry, extracellular single cells could be easily gated for size by scattering. However, we observed that sometimes bacteria appear bound to immune cells even though they have not been internalized, resulting in a single event that can hardly be identified through scattering or in the brightfield image of imaging-based flow cytometry. CouCyCF_3_ **1b** allows for the visualization of these extracellular bacterial cells (Figure 4A, Figure S19), providing an additional feature to track pre-infection events.

**Figure 4:**
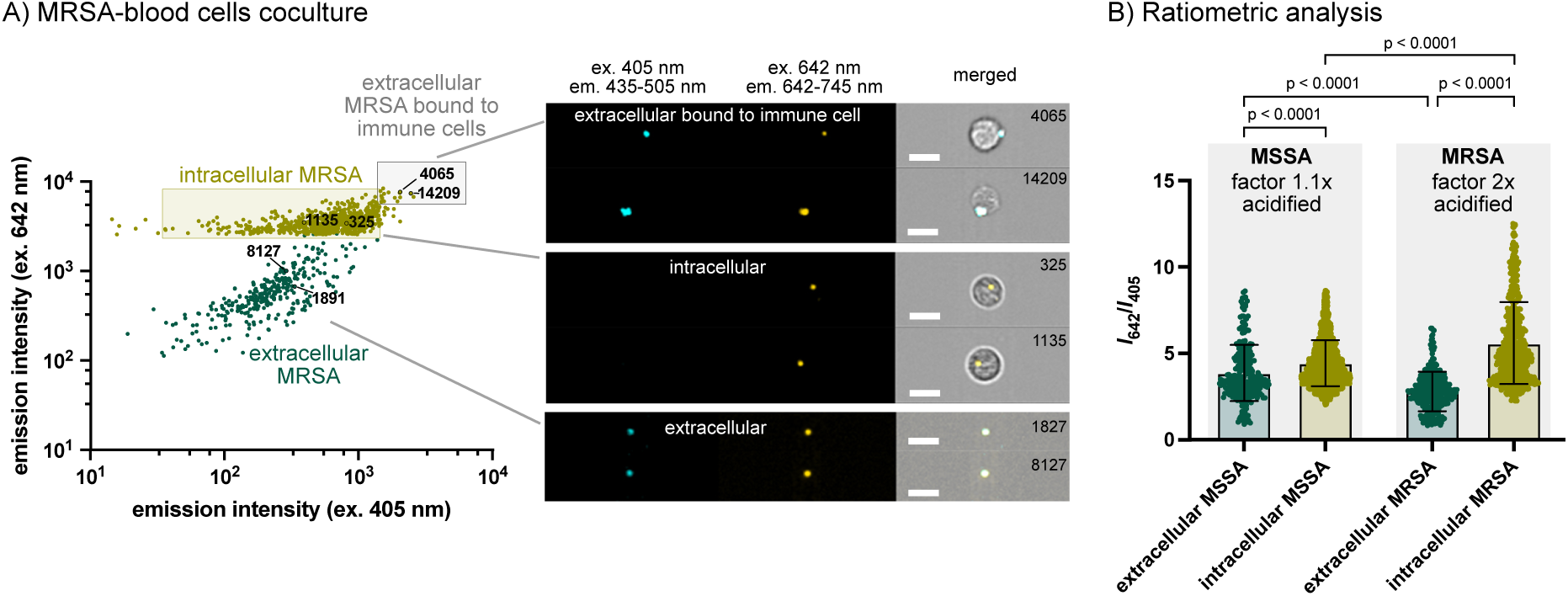
Phagocytosis study with clinical isolates of *S. aureus* and blood cells. A) Scatter plot and images of extra- and intracellular MRSA cells stained with CouCyCF3 **1b** (1 µM) 10 min post-infection. Internalized bacteria (yellow; events 1135 and 325) in blood cells undergo phagocytosis, represented as a population shift to higher red emission intensities compared to the extracellular bacteria population (green; events 1827 and 8127). Pre-infection events of extracellular bacteria bound to immune cells appeared with higher blue and red emission intensities (events 4065 and 14209). Merged images display the overlay of the brightfield and emission channels. Scale bar, 7 µm. B) Quantification of internal pH based on the *I*red/*I*blue emission intensities. Graph represents single cell intensities (scatter plot) and means (bar plot) with SD (error bar) for extracellular MSSA (mean = 3.9 ± 1.6, *N* = 270), intracellular MSSA (mean = 4.4 ± 1.3, *N* = 2246), extracellular MRSA (mean = 2.8 ± 1.1, *N* = 325), and intracellular MRSA (mean = 5.6 ± 1.1, *N* = 553). Statistical significance was evaluated using the Kruskal-Wallis test (multiple comparisons). Outliers were identified and removed using the ROUT method (Q = 1%) with 5%, 6%, 1%, and 12% outliers (from left to right).

Our quantification of the *I*_red_/*I*_blue_ ratio showed that extracellular MSSA cells have a more acidic intracellular pH compared to MRSA (Figure 4B). Interestingly, this trend was reversed upon phagocytosis: intracellular MSSA cells were less acidified, showing only a 1.1-fold decrease in pH, whereas MRSA cells experienced a 2-fold acidification. The data suggest that MSSA exhibit increased tolerance to acidic stress during phagocytosis. This resilience might be attributed to the pre-activation of the acid tolerance response (ATR), a cellular defense mechanism initiated by exposure to mildly acidic conditions in either extracellular or intracellular environments.^22,25^ The reduced cytoplasmic pH of extracellular MSSA might have induced ATR, enabling these cells to better tolerate the harsh acidic environment inside phagolysosomes. This tolerance is likely based on pH-dependent gene regulation; mildly acidic pH modulates a large set of staphylococcal genes, including virulence, resulting in a cellular remodeling that adapts bacterial phenotypes to pH-variable environments.^85^ Our observations align with previous reports demonstrating that ATR enhances bacterial survival under lethal acidic conditions, including the strong acidic environment within macrophages.^25,47,86^ Previous research has shown that stress-resilient phenotypes that evolve in persister cells have a lower cytoplasmic pH.^31^ Similarly, our data indicate that more acidified MSSA are less affected by acid stress during phagocytosis, suggesting that low intracellular pH may serve as a predictive marker to help identify tolerant cells before stress exposure.

It is important to note that for this proof-of-concept study, we report population-averaged *I*_red_/*I*_blue_ ratios. However, CouCyCF_3_ **1b** allows for visualization of single-cell traits, providing the potential to analyze heterogeneity in stress adaptation at the individual cell level. Overall, these results demonstrate that CouCyCF_3_ **1b** can be used to track phagocytosis and visualize differences in the physiological state of clinical *S. aureus* strains in complex biological samples.

## Conclusion

We report a ratiometric fluorescent sensor specifically designed to study the intracellular pH of bacteria. The CouCy scaffold reacts fast and reversibly with OH^-^ ions and induces a ratiometric emission change that can be quantified as *I*_red_/*I*_blue_. The sensitivity of the scaffold was successfully adjusted by introducing EWG to the indoleninium core, resulting in CouCyCF_3_ **1b** having a biologically relevant, large dynamic sensing range (pH 5.0-7.5). We demonstrated live-cell pH sensing in *E. coli*, *S. epidermidis*, and clinical isolates of *S. aureus* using fluorescence microscopy and flow cytometry. Our probe has a higher sensitivity than the commercially available small-molecule BCECF-AM **S1** or the protein sensor pHluorin. This feature enabled the visualization of pH-sensitive *E. coli* cells, highlighting the ability to visualize single-cell phenotypes in real-time. Additionally, the phagocytosis of clinical *S. aureus* strains in THP-1 monocytes or blood samples could be monitored.

Our studies show that CouCyCF_3_ **1b** is a general pH sensing probe. Despite the high uptake and accumulation in bacteria, further improvements could be made regarding its retention in bacteria. To reduce efflux over time, functional groups for intracellular trapping could be installed, such as an isothiocyanate or an *N*-hydroxysuccinimide (NHS) ester, as used in many commercially available probes. CouCy probes are an excellent scaffold for bacterial imaging and a promising tool for detecting phenotypic heterogeneity, which, with further fine-tuning, might allow unraveling the physiological state of resistant or persistent bacteria of clinical relevance.

## Associated content

### Supporting Information

A supporting information file is available free of charge. The contents of this file include experimental details, supplementary figures, and ^1^H and ^13^C NMR spectra (PDF).

### Author contributions

D. K. and P. R.-F. conceived of the study. D. K. performed all experiments and analyzed all the data, except for those involving clinical isolates, which were performed in collaboration with A. I. The work with clinical strains was supervised by N. K. The manuscript was written by D. K. and P. R.-F. with input from A. I. and N. K.

### Notes

The collection, pseudonymization, and usage of clinical samples used in this project was carried out with the appropriate ethics approval within the NCCR AntiResist project (2020-02588). The authors declare no competing interests. All the raw data from this paper are available on Zenodo at DOI: 10.5281/zenodo.17034283.

## Acknowledgments

This work was funded by the Swiss National Science Foundation (NCCR AntiResist, grant no. 180541). We thank Dr. Alina Tirla for providing some chemical precursors and Emre Demirbilek for support with experiments involving clinical isolates. Flow Cytometry was performed at the Cytometry Facilities at the University of Zurich and the University of Basel.

## References

(1) Murray, C. J.; Ikuta, K. S.; Sharara, F.; Swetschinski, L.; Robles Aguilar, G.; Gray, A.; Han, C.; Bisignano, C.; Rao, P.; Wool, E.; Johnson, S. C.; Browne, A. J.; Chipeta, M. G.; Fell, F.; Hackett, S.; Haines-Woodhouse, G.; Kashef Hamadani, B. H.; Kumaran, E. A. P.; McManigal, B.; Agarwal, R.; Akech, S.; Albertson, S.; Amuasi, J.; Andrews, J.; Aravkin, A.; Ashley, E.; Bailey, F.; Baker, S.; Basnyat, B.; Bekker, A.; Bender, R.; Bethou, A.; Bielicki, J.; Boonkasidecha, S.; Bukosia, J.; Carvalheiro, C.; Castañeda-Orjuela, C.; Chansamouth, V.; Chaurasia, S.; Chiurchiù, S.; Chowdhury, F.; Cook, A. J.; Cooper, B.; Cressey, T. R.; Criollo-Mora, E.; Cunningham, M.; Darboe, S.; Day, N. P. J.; De Luca, M.; Dokova, K.; Dramowski, A.; Dunachie, S. J.; Eckmanns, T.; Eibach, D.; Emami, A.; Feasey, N.; Fisher-Pearson, N.; Forrest, K.; Garrett, D.; Gastmeier, P.; Giref, A. Z.; Greer, R. C.; Gupta, V.; Haller, S.; Haselbeck, A.; Hay, S. I.; Holm, M.; Hopkins, S.; Iregbu, K. C.; Jacobs, J.; Jarovsky, D.; Javanmardi, F.; Khorana, M.; Kissoon, N.; Kobeissi, E.; Kostyanev, T.; Krapp, F.; Krumkamp, R.; Kumar, A.; Kyu, H. H.; Lim, C.; Limmathurotsakul, D.; Loftus, M. J.; Lunn, M.; Ma, J.; Mturi, N.; Munera-Huertas, T.; Musicha, P.; Mussi-Pinhata, M. M.; Nakamura, T.; Nanavati, R.; Nangia, S.; Newton, P.; Ngoun, C.; Novotney, A.; Nwakanma, D.; Obiero, C. W.; Olivas-Martinez, A.; Olliaro, P.; Ooko, E.; Ortiz-Brizuela, E.; Peleg, A. Y.; Perrone, C.; Plakkal, N.; Ponce-de-Leon, A.; Raad, M.; Ramdin, T.; Riddell, A.; Roberts, T.; Robotham, J. V.; Roca, A.; Rudd, K. E.; Russell, N.; Schnall, J.; Scott, J. A. G.; Shivamallappa, M.; Sifuentes-Osornio, J.; Steenkeste, N.; Stewardson, A. J.; Stoeva, T.; Tasak, N.; Thaiprakong, A.; Thwaites, G.; Turner, C.; Turner, P.; van Doorn, H. R.; Velaphi, S.; Vongpradith, A.; Vu, H.; Walsh, T.; Waner, S.; Wangrangsimakul, T.; Wozniak, T.; Zheng, P.; Sartorius, B.; Lopez, A. D.; Stergachis, A.; Moore, C.; Dolecek, C.; Naghavi, M. Global Burden of Bacterial Antimicrobial Resistance in 2019: A Systematic Analysis. The Lancet 2022, 399 (10325), 629–655. 10.1016/S0140-6736(21)02724-0.

(2) Bigger, J. W. Treatment of Staphylococcal Infections with Penicillin by Intermittent Sterilisation. Lancet 1944, 244 (6320), 497–500. 10.1016/S0140-6736(00)74210-3.

(3) Hughes, D.; Andersson, D. I. Environmental and Genetic Modulation of the Phenotypic Expression of Antibiotic Resistance. FEMS Microbiology Reviews 2017, 41 (3), 374–391. 10.1093/femsre/fux004.

(4) Huemer, M.; Mairpady Shambat, S.; Brugger, S. D.; Zinkernagel, A. S. Antibiotic Resistance and Persistence-Implications for Human Health and Treatment Perspectives. EMBO Rep. 2020, 21 (12), e51034. 10.15252/embr.202051034.

(5) Şimşek, E.; Kim, M. The Emergence of Metabolic Heterogeneity and Diverse Growth Responses in Isogenic Bacterial Cells. ISME J. 2018, 12 (5), 1199–1209. 10.1038/s41396-017-0036-2.

(6) Nikolic, N.; Barner, T.; Ackermann, M. Analysis of Fluorescent Reporters Indicates Heterogeneity in Glucose Uptake and Utilization in Clonal Bacterial Populations. BMC Microbiol. 2013, 13 (1), 258. 10.1186/1471-2180-13-258.

(7) Kotte, O.; Volkmer, B.; Radzikowski, J. L.; Heinemann, M. Phenotypic Bistability in Escherichia Coli’s Central Carbon Metabolism. Mol. Syst. Biol. 2014, 10 (7), 736. 10.15252/msb.20135022.

(8) Balaban, N. Q.; Merrin, J.; Chait, R.; Kowalik, L.; Leibler, S. Bacterial Persistence as a Phenotypic Switch. Science 2004, 305 (5690), 1622–1625. 10.1126/science.1099390.

(9) Wilmaerts, D.; Windels, E. M.; Verstraeten, N.; Michiels, J. General Mechanisms Leading to Persister Formation and Awakening. Trends Genet. 2019, 35 (6), 401–411. 10.1016/j.tig.2019.03.007.

(10) Mordukhova, E. A.; Pan, J.-G. Stabilization of Homoserine-O-Succinyltransferase (MetA) Decreases the Frequency of Persisters in Escherichia Coli under Stressful Conditions. PLOS ONE 2014, 9 (10), e110504. 10.1371/journal.pone.0110504.

(11) Murakami, K.; Ono, T.; Viducic, D.; Kayama, S.; Mori, M.; Hirota, K.; Nemoto, K.; Miyake, Y. Role for *rpoS* Gene of *Pseudomonas Aeruginosa* in Antibiotic Tolerance. FEMS Microbiol. Lett. 2005, 242 (1), 161–167. 10.1016/j.femsle.2004.11.005.

(12) Sollier, J.; Basler, M.; Broz, P.; Dittrich, P. S.; Drescher, K.; Egli, A.; Harms, A.; Hierlemann, A.; Hiller, S.; King, C. G.; McKinney, J. D.; Moran-Gilad, J.; Neher, R. A.; Page, M. G. P.; Panke, S.; Persat, A.; Picotti, P.; Rentsch, K. M.; Rivera-Fuentes, P.; Sauer, U.; Stolz, D.; Tschudin-Sutter, S.; van Delden, C.; van Nimwegen, E.; Veening, J.-W.; Zampieri, M.; Zinkernagel, A. S.; Khanna, N.; Bumann, D.; Jenal, U.; Dehio, C. Revitalizing Antibiotic Discovery and Development through in Vitro Modelling of In-Patient Conditions. Nat. Microbiol. 2024, 9 (1), 1–3. 10.1038/s41564-023-01566-w.

(13) Spratt, M. R.; Lane, K. Navigating Environmental Transitions: The Role of Phenotypic Variation in Bacterial Responses. mBio 2022, 13 (6), e02212–22. 10.1128/mbio.02212-22.

(14) Brauner, A.; Fridman, O.; Gefen, O.; Balaban, N. Q. Distinguishing between Resistance, Tolerance and Persistence to Antibiotic Treatment. Nat. Rev. Microbiol. 2016, 14 (5), 320–330. 10.1038/nrmicro.2016.34.

(15) Gefen, O.; Balaban, N. Q. The Importance of Being Persistent: Heterogeneity of Bacterial Populations under Antibiotic Stress. FEMS Microbiol. Rev. 2009, 33 (4), 704–717. 10.1111/j.1574-6976.2008.00156.x.

(16) Gefen, O.; Chekol, B.; Strahilevitz, J.; Balaban, N. Q. TDtest: Easy Detection of Bacterial Tolerance and Persistence in Clinical Isolates by a Modified Disk-Diffusion Assay. Sci Rep 2017, 7 (1), 41284. 10.1038/srep41284.

(17) Bär, J.; Boumasmoud, M.; Kouyos, R. D.; Zinkernagel, A. S.; Vulin, C. Efficient Microbial Colony Growth Dynamics Quantification with ColTapp, an Automated Image Analysis Application. Sci. Rep. 2020, 10 (1), 16084. 10.1038/s41598-020-72979-4.

(18) Levin-Reisman, I.; Gefen, O.; Fridman, O.; Ronin, I.; Shwa, D.; Sheftel, H.; Balaban, N. Q. Automated Imaging with ScanLag Reveals Previously Undetectable Bacterial Growth Phenotypes. Nat. Methods 2010, 7 (9), 737–739. 10.1038/nmeth.1485.

(19) Specht, E. A.; Braselmann, E.; Palmer, A. E. A Critical and Comparative Review of Fluorescent Tools for Live-Cell Imaging. Annu. Rev. of Physiol. 2017, 79, 93–117. 10.1146/annurev-physiol-022516-034055.

(20) Hare, P. J.; LaGree, T. J.; Byrd, B. A.; DeMarco, A. M.; Mok, W. W. K. Single-Cell Technologies to Study Phenotypic Heterogeneity and Bacterial Persisters. Microorganisms 2021, 9 (11), 2277. 10.3390/microorganisms9112277.

(21) Krulwich, T. A.; Sachs, G.; Padan, E. Molecular Aspects of Bacterial pH Sensing and Homeostasis. Nat. Rev. Microbiol. 2011, 9 (5), 330–343. 10.1038/nrmicro2549.

(22) Lund, P.; Tramonti, A.; De Biase, D. Coping with Low pH: Molecular Strategies in Neutralophilic Bacteria. FEMS Microbiol. Rev. 2014, 38 (6), 1091–1125. 10.1111/1574-6976.12076.

(23) Choi, J.; Groisman, E. A. Acidic pH Sensing in the Bacterial Cytoplasm Is Required for Salmonella Virulence. Mol. Microbiol. 2016, 101 (6), 1024–1038. 10.1111/mmi.13439.

(24) Leimer, N.; Rachmühl, C.; Palheiros Marques, M.; Bahlmann, A. S.; Furrer, A.; Eichenseher, F.; Seidl, K.; Matt, U.; Loessner, M. J.; Schuepbach, R. A.; Zinkernagel, A. S. Nonstable Staphylococcus Aureus Small-Colony Variants Are Induced by Low pH and Sensitized to Antimicrobial Therapy by Phagolysosomal Alkalinization. J. Infect. Dis. 2016, 213 (2), 305–313. 10.1093/infdis/jiv388.

(25) O’Sullivan, E.; Condon, S. Intracellular pH Is a Major Factor in the Induction of Tolerance to Acid and Other Stresses in Lactococcus Lactis. Appl. Environ. Microbiol. 1997, 63 (11), 4210–4215. 10.1128/aem.63.11.4210-4215.1997.

(26) Padan, E.; Bibi, E.; Ito, M.; Krulwich, T. A. Alkaline pH Homeostasis in Bacteria: New Insights. Biochim. Biophys. Acta. 2005, 1717 (2), 67–88. 10.1016/j.bbamem.2005.09.010.

(27) Padan, E.; Zilberstein, D.; Schuldiner, S. pH Homesstasis in Bacteria. *Biochim. Biophys. Acta*, Biomembr. 1981, 650 (2), 151–166. 10.1016/0304-4157(81)90004-6.

(28) Slonczewski, J. L.; Rosen, B. P.; Alger, J. R.; Macnab, R. M. pH Homeostasis in Escherichia Coli: Measurement by 31P Nuclear Magnetic Resonance of Methylphosphonate and Phosphate. PNAS 1981, 78 (10), 6271–6275. 10.1073/pnas.78.10.6271.

(29) Sedlyarov, V.; Eichner, R.; Girardi, E.; Essletzbichler, P.; Goldmann, U.; Nunes-Hasler, P.; Srndic, I.; Moskovskich, A.; Heinz, L. X.; Kartnig, F.; Bigenzahn, J. W.; Rebsamen, M.; Kovarik, P.; Demaurex, N.; Superti-Furga, G. The Bicarbonate Transporter SLC4A7 Plays a Key Role in Macrophage Phagosome Acidification. Cell Host Microbe 2018, 23 (6), 766–774.e5. 10.1016/j.chom.2018.04.013.

(30) McKinney, J. D.; zu Bentrup, K. H.; Muñoz-Elías, E. J.; Miczak, A.; Chen, B.; Chan, W.-T.; Swenson, D.; Sacchettini, J. C.; Jacobs, W. R.; Russell, D. G. Persistence of Mycobacterium Tuberculosis in Macrophages and Mice Requires the Glyoxylate Shunt Enzyme Isocitrate Lyase. Nature 2000, 406 (6797), 735–738. 10.1038/35021074.

(31) Goode, O.; Smith, A.; Zarkan, A.; Cama, J.; Invergo, B. M.; Belgami, D.; Caño-Muñiz, S.; Metz, J.; O’Neill, P.; Jeffries, A.; Norville, I. H.; David, J.; Summers, D.; Pagliara, S. Persister Escherichia Coli Cells Have a Lower Intracellular pH than Susceptible Cells but Maintain Their pH in Response to Antibiotic Treatment. mBio 2021, 12 (4), e00909–21. 10.1128/mbio.00909-21.

(32) Miesenböck, G.; De Angelis, D. A.; Rothman, J. E. Visualizing Secretion and Synaptic Transmission with pH-Sensitive Green Fluorescent Proteins. Nature 1998, 394, 192–195. 10.1038/28190.

(33) Sankaranarayanan, S.; Angelis, D. D.; Rothman, J. E.; Ryan, T. A. The Use of pHluorins for Optical Measurements of Presynaptic Activity. Biophys. J. 2000, 79 (4), 2199–2208. 10.1016/S0006-3495(00)76468-X.

(34) Zarkan, A.; Caño-Muñiz, S.; Zhu, J.; Al Nahas, K.; Cama, J.; Keyser, U. F.; Summers, D. K. Indole Pulse Signalling Regulates the Cytoplasmic pH of E. Coli in a Memory-Like Manner. Sci. Rep. 2019, 9 (1), 3868. 10.1038/s41598-019-40560-3.

(35) Tomasek, K.; Bergmiller, T.; Guet, C. C. Lack of Cations in Flow Cytometry Buffers Affect Fluorescence Signals by Reducing Membrane Stability and Viability of Escherichia Coli Strains. J. Biotech. 2018, 268, 40–52. 10.1016/j.jbiotec.2018.01.008.

(36) Craggs, T. D. Green Fluorescent Protein: Structure, Folding and Chromophore Maturation. Chem. Soc. Rev. 2009, 38 (10), 2865–2875. 10.1039/B903641P.

(37) Arce-Rodríguez, A.; Volke, D. C.; Bense, S.; Häussler, S.; Nikel, P. I. Non-Invasive, Ratiometric Determination of Intracellular pH in Pseudomonas Species Using a Novel Genetically Encoded Indicator. Microb. Biotechnol. 2019, 12 (4), 799–813. 10.1111/1751-7915.13439.

(38) Olsen, K. N.; Budde, B. B.; Siegumfeldt, H.; Rechinger, K. B.; Jakobsen, M.; Ingmer, H. Noninvasive Measurement of Bacterial Intracellular pH on a Single-Cell Level with Green Fluorescent Protein and Fluorescence Ratio Imaging Microscopy. Appl. Environ. Microbiol. 2002, 68 (8), 4145–4147. 10.1128/AEM.68.8.4145-4147.2002.

(39) van Beilen, J. W. A.; Brul, S. Compartment-Specific pH Monitoring in Bacillus Subtilis Using Fluorescent Sensor Proteins: A Tool to Analyze the Antibacterial Effect of Weak Organic Acids. Front. Microbiol. 2013, 4. 10.3389/fmicb.2013.00157.

(40) Jantarug, K.; Tripathi, V.; Morin, B.; Iizuka, A.; Kuehl, R.; Morgenstern, M.; Clauss, M.; Khanna, N.; Bumann, D.; Rivera-Fuentes, P. A Far-Red Fluorescent Probe to Visualize Gram-Positive Bacteria in Patient Samples. ACS Infect. Dis. 2024, 10 (5), 1545–1551. 10.1021/acsinfecdis.4c00060.

(41) Ropponen, H.-K.; Richter, R.; Hirsch, A. K. H.; Lehr, C.-M. Mastering the Gram-Negative Bacterial Barrier – Chemical Approaches to Increase Bacterial Bioavailability of Antibiotics. Adv. Drug Deliv. Rev. 2021, 172, 339–360. 10.1016/j.addr.2021.02.014.

(42) Richter, M. F.; Hergenrother, P. J. The Challenge of Converting Gram-Positive-Only Compounds into Broad-Spectrum Antibiotics. Ann. N. Y. Acad. Sci. 2019, 1435, 18–38. 10.1111/nyas.13598.

(43) Wunsch, C. M.; Lewis, J. P. Porphyromonas Gingivalis as a Model Organism for Assessing Interaction of Anaerobic Bacteria with Host Cells. J. Vis. Exp. 2015, No. 106, e53408. 10.3791/53408.

(44) Hayes, E.; Murphy, M. P.; Pohl, K.; Browne, N.; McQuillan, K.; Saw, L. E.; Foley, C.; Gargoum, F.; McElvaney, O. J.; Hawkins, P.; Gunaratnam, C.; McElvaney, N. G.; Reeves, E. P. Altered Degranulation and pH of Neutrophil Phagosomes Impacts Antimicrobial Efficiency in Cystic Fibrosis. Front. Immunol. 2020, 11. 10.3389/fimmu.2020.600033.

(45) Vergne, I.; Constant, P.; Lanéelle, G. Phagosomal pH Determination by Dual Fluorescence Flow Cytometry. Analytical Biochemistry 1998, 255 (1), 127–132. 10.1006/abio.1997.2466.

(46) Liu, J.; Diwu, Z.; Leung, W.-Y. Synthesis and Photophysical Properties of New Fluorinated Benzo[*c*]Xanthene Dyes as Intracellular pH Indicators. Bioorg. Med. Chem. Lett. 2001, 11 (22), 2903–2905. 10.1016/S0960-894X(01)00595-9.

(47) Schneider, B.; Gross, R.; Haas, A. Phagosome Acidification Has Opposite Effects on Intracellular Survival of Bordetella Pertussis andB. Bronchiseptica. Infect. Immun. 2000, 68 (12), 7039–7048. 10.1128/iai.68.12.7039-7048.2000.

(48) Porte, F.; Liautard, J.-P.; Köhler, S. Early Acidification of Phagosomes Containing Brucella Suis Is Essential for Intracellular Survival in Murine Macrophages. Infect. Immun. 1999, 67 (8), 4041–4047. 10.1128/iai.67.8.4041-4047.1999.

(49) Hunter, R. C.; Beveridge, T. J. Application of a pH-Sensitive Fluoroprobe (C-SNARF-4) for pH Microenvironment Analysis in Pseudomonas Aeruginosa Biofilms. Appl. Environ. Microbiol. 2005, 71 (5), 2501–2510. 10.1128/AEM.71.5.2501-2510.2005.

(50) Schlafer, S.; Garcia, J. E.; Greve, M.; Raarup, M. K.; Nyvad, B.; Dige, I. Ratiometric Imaging of Extracellular pH in Bacterial Biofilms with C-SNARF-4. Appl. Environ. Microbiol. 2015, 81 (4), 1267–1273. 10.1128/AEM.02831-14.

(51) Richter, M. F.; Drown, B. S.; Riley, A. P.; Garcia, A.; Shirai, T.; Svec, R. L.; Hergenrother, P. J. Predictive Compound Accumulation Rules Yield a Broad-Spectrum Antibiotic. Nature 2017, 545 (7654), 299–304. 10.1038/nature22308.

(52) Zhang, Y.; Chen, Y.; Fang, H.; Wang, Y.; Li, S.; Yuan, H.; Yao, S.; Qin, S.; He, W.; Guo, Z. A Ratiometric pH Probe for Acidification Tracking in Dysfunctional Mitochondria and Tumour Tissue in Vivo. J. Mater. Chem. B 2022, 10 (28), 5422– 5429. 10.1039/D2TB00553K.

(53) Chen, S.; Hong, Y.; Liu, Y.; Liu, J.; Leung, C. W. T.; Li, M.; Kwok, R. T. K.; Zhao, E.; Lam, J. W. Y.; Yu, Y.; Tang, B. Z. Full-Range Intracellular pH Sensing by an Aggregation-Induced Emission-Active Two-Channel Ratiometric Fluorogen. J. Am. Chem. Soc. 2013, 135 (13), 4926–4929. 10.1021/ja400337p.

(54) Peng, J.; Chen, H.; Sun, M.; Yu, H.; Hou, J.; Wang, S. A Benzooxazine-Based Ratiometric Fluorescent Probe for pH Imaging in Living Cells and Bacteria. Sens. Actuators C Chem. 2021, 335, 129711. 10.1016/j.snb.2021.129711.

(55) Tirla, A.; Rivera-Fuentes, P. Development of a Photoactivatable Phosphine Probe for Induction of Intracellular Reductive Stress with Single-Cell Precision. Angew. Chem. Int. Ed. 2016, 55 (47), 14709–14712. 10.1002/anie.201608779.

(56) Ho, T.-L. Hard Soft Acids Bases (HSAB) Principle and Organic Chemistry. Chem. Rev. 1975, 75 (1), 1–20. 10.1021/cr60293a001.

(57) Liang, Z.; Sun, Y.; Duan, R.; Yang, R.; Qu, L.; Zhang, K.; Li, Z. Low Polarity-Triggered Basic Hydrolysis of Coumarin as an AND Logic Gate for Broad-Spectrum Cancer Diagnosis. Anal. Chem. 2021, 93 (36), 12434–12440. 10.1021/acs.analchem.1c02591.

(58) Peng, M.-J.; Guo, Y.; Yang, X.-F.; Suzenet, F.; Li, J.; Li, C.-W.; Duan, Y.-W. Coumarin–Hemicyanine Conjugates as Novel Reaction-Based Sensors for Cyanide Detection: Convenient Synthesis and ICT Mechanism. RSC Adv. 2014, 4 (37), 19077–19085. 10.1039/C4RA01598C.

(59) Lv, X.; Liu, J.; Liu, Y.; Zhao, Y.; Sun, Y.-Q.; Wang, P.; Guo, W. Ratiometric Fluorescence Detection of Cyanide Based on a Hybrid Coumarin–Hemicyanine Dye: The Large Emission Shift and the High Selectivity. Chem. Commun. 2011, 47 (48), 12843–12845. 10.1039/C1CC15721C.

(60) Chen, Y.; Zhu, C.; Yang, Z.; Chen, J.; He, Y.; Jiao, Y.; He, W.; Qiu, L.; Cen, J.; Guo, Z. A Ratiometric Fluorescent Probe for Rapid Detection of Hydrogen Sulfide in Mitochondria. Angew. Chem. Int. Ed. 2013, 52 (6), 1688–1691. 10.1002/anie.201207701.

(61) Wang, B.; Jiang, N.; Sun, W.; Wang, Q.; Zheng, G. A Ratiometric Fluorescence Probe for Detection of Hydrogen Sulfide in Cells. RSC Adv. 2016, 6 (43), 36906– 36909. 10.1039/C6RA02579J.

(62) Yin, G.; Niu, T.; Gan, Y.; Yu, T.; Yin, P.; Chen, H.; Zhang, Y.; Li, H.; Yao, S. A Multi-Signal Fluorescent Probe with Multiple Binding Sites for Simultaneous Sensing of Cysteine, Homocysteine, and Glutathione. Angew. Chem. Int. Ed. 2018, 57 (18), 4991–4994. 10.1002/anie.201800485.

(63) Xiong, K.; Huo, F.; Chao, J.; Zhang, Y.; Yin, C. Colorimetric and NIR Fluorescence Probe with Multiple Binding Sites for Distinguishing Detection of Cys/Hcy and GSH in Vivo. Anal. Chem. 2019, 91 (2), 1472–1478. 10.1021/acs.analchem.8b04485.

(64) Ryall, B.; Davies, J. C.; Wilson, R.; Shoemark, A.; Williams, H. D. Pseudomonas Aeruginosa, Cyanide Accumulation and Lung Function in CF and Non-CF Bronchiectasis Patients. Eur. Respir. J. 2008, 32 (3), 740–747. 10.1183/09031936.00159607.

(65) Newton, G. L.; Arnold, K.; Price, M. S.; Sherrill, C.; Delcardayre, S. B.; Aharonowitz, Y.; Cohen, G.; Davies, J.; Fahey, R. C.; Davis, C. Distribution of Thiols in Microorganisms: Mycothiol Is a Major Thiol in Most Actinomycetes. J. Bacteriol. 1996, 178 (7), 1990–1995. 10.1128/jb.178.7.1990-1995.1996.

(66) Ferreira, R. J.; Kasson, P. M. Antibiotic Uptake across Gram-Negative Outer Membranes: Better Predictions towards Better Antibiotics. ACS Infect. Dis. 2019, 5, 2096–2104. 10.1021/acsinfecdis.9b00201.

(67) Madhu, M.; Santhoshkumar, S.; Tseng, W.-B.; Tseng, W.-L. Maximizing Analytical Precision: Exploring the Advantages of Ratiometric Strategy in Fluorescence, Raman, Electrochemical, and Mass Spectrometry Detection. Front. Anal. Sci. 2023, 3, 1–14. 10.3389/frans.2023.1258558.

(68) Checa, M.; Millan-Solsona, R.; Blanco, N.; Torrents, E.; Fabregas, R.; Gomila, G. Mapping the Dielectric Constant of a Single Bacterial Cell at the Nanoscale with Scanning Dielectric Force Volume Microscopy. Nanoscale 2019, 11 (43), 20809–20819. 10.1039/C9NR07659J.

(69) Yoon, S. A.; Cha, S. H.; Jun, S. W.; Park, S. J.; Park, J.-Y.; Lee, S.; Kim, H. S.; Ahn, Y. H. Identifying Different Types of Microorganisms with Terahertz Spectroscopy. Biomed. Opt. Express 2019, 11 (1), 406–416. 10.1364/BOE.376584.

(70) Esteban-Ferrer, D.; Edwards, M. A.; Fumagalli, L.; Juárez, A.; Gomila, G. Electric Polarization Properties of Single Bacteria Measured with Electrostatic Force Microscopy. ACS Nano 2014, 8 (10), 9843–9849. 10.1021/nn5041476.

(71) Plášek, J.; Babuka, D.; Hoefer, M. H+ Translocation by Weak Acid Uncouplers Is Independent of H+ Electrochemical Gradient. J. Bioenerg. Biomembr. 2017, 49 (5), 391–397. 10.1007/s10863-017-9724-x.

(72) Rink, T. J.; Tsien, R. Y.; Pozzan, T. Cytoplasmic pH and Free Mg2+ in Lymphocytes. JCB 1982, 95 (1), 189–196. 10.1083/jcb.95.1.189.

(73) Gelman-Zhornitsky, E.; Deutsch, M.; Tirosh, R.; Yishay, Y.; Weinreb, A.; Shapiro, H. M. 2’,7’-Bis-(Carboxyethyl)-5-(6’)-Caroboxyfluorescein (BCECF) as a Probe for Intracellular Fluorescence Polarization Measurements. JBO 1997, 2 (2), 186–194. 10.1117/12.251737.

(74) Klonis, N.; Sawyer, W. H. Spectral Properties of the Prototropic Forms of Fluorescein in Aqueous Solution. J. Fluoresc. 1996, 6 (3), 147–157. 10.1007/BF00732054.

(75) Brown, J. L.; Ross, T.; McMeekin, T. A.; Nichols, P. D. Acid Habituation of *Escherichia Coli* and the Potential Role of Cyclopropane Fatty Acids in Low pH Tolerance. Int. J. Food Microbiol. 1997, 37 (2), 163–173. 10.1016/S0168-1605(97)00068-8.

(76) Chang, Y.-Y.; Cronan, J. E. Membrane Cyclopropane Fatty Acid Content Is a Major Factor in Acid Resistance of Escherichia Coli. Mol. Microbiol. 1999, 33 (2), 249–259. 10.1046/j.1365-2958.1999.01456.x.

(77) Shabala, L.; Ross, T. Cyclopropane Fatty Acids Improve *Escherichia Coli* Survival in Acidified Minimal Media by Reducing Membrane Permeability to H+ and Enhanced Ability to Extrude H+. Microbiol. Res. 2008, 159 (6), 458–461. 10.1016/j.resmic.2008.04.011.

(78) Wang, A.-Y.; Cronan Jr, J. E. The Growth Phase-Dependent Synthesis of Cyclopropane Fatty Acids in Escherichia Coli Is the Result of an RpoS(KatF)-Dependent Promoter plus Enzyme Instability. Mol. Microbiol. 1994, 11 (6), 1009–1017. 10.1111/j.1365-2958.1994.tb00379.x.

(79) Datsenko, K. A.; Wanner, B. L. One-Step Inactivation of Chromosomal Genes in Escherichia Coli K-12 Using PCR Products. Proc. Natl. Acad. Sci. U.S.A 2000, 97 (12), 6640–6645. 10.1073/pnas.120163297.

(80) Baba, T.; Ara, T.; Hasegawa, M.; Takai, Y.; Okumura, Y.; Baba, M.; Datsenko, K. A.; Tomita, M.; Wanner, B. L.; Mori, H. Construction of Escherichia Coli K-12 in-Frame, Single-Gene Knockout Mutants: The Keio Collection. Mol. Syst. Biol. 2006, 2, 2006.0008. 10.1038/msb4100050.

(81) Maiti, A.; Kumar, A.; Daschakraborty, S. How Do Cyclopropane Fatty Acids Protect the Cell Membrane of Escherichia Coli in Cold Shock? J. Phys. Chem. B 2023, 127 (7), 1607–1617. 10.1021/acs.jpcb.3c00541.

(82) Richards, T. A.; Massana, R.; Pagliara, S.; Hall, N. Single Cell Ecology. Philos. Trans. R. Soc. Lond. B Biol. Sci. 2019, 374 (1786), 20190076. 10.1098/rstb.2019.0076.

(83) Lenzo, J. C.; O’Brien-Simpson, N. M.; Cecil, J.; Holden, J. A.; Reynolds, E. C. Determination of Active Phagocytosis of Unopsonized Porphyromonas Gingivalis by Macrophages and Neutrophils Using the pH-Sensitive Fluorescent Dye pHrodo. Infect. Immun. 2016, 84 (6), 1753–1760. 10.1128/iai.01482-15.

(84) Ogawa, M.; Kosaka, N.; S. Regino, C. A.; Mitsunaga, M.; L. Choyke, P.; Kobayashi, H. High Sensitivity Detection of Cancer in Vivo Using a Dual-Controlled Activation Fluorescent Imaging Probe Based on H-Dimer Formation and pH Activation. Mol. Biosyst. 2010, 6 (5), 888–893. 10.1039/B917876G.

(85) Weinrick, B.; Dunman, P. M.; McAleese, F.; Murphy, E.; Projan, S. J.; Fang, Y.; Novick, R. P. Effect of Mild Acid on Gene Expression in Staphylococcus Aureus. J. Bacteriol. 2004, 186 (24), 8407–8423. 10.1128/jb.186.24.8407-8423.2004.

(86) Wilmes-Riesenberg, M. R.; Bearson, B.; Foster, J. W.; Curtis, R. Role of the Acid Tolerance Response in Virulence of Salmonella Typhimurium. Infect. Immun. 1996, 64 (4), 1085–1092. 10.1128/iai.64.4.1085-1092.1996.

