## Supplementary Information for "A ratiometric pH sensor for Gram-positive and Gram-negative bacteria"

#### General Remarks

All starting materials were purchased from commercial sources and used without further purification. Solvents were procured from Sigma-Aldrich or Thommen-Furler. Reactions were monitored by mass spectrometry and thin layer chromatography (TLC, silica gel 60 $F_{254}$, Sigma-Aldrich) using UV light at 254 nm and 366 nm for visualization. Purification of reaction products was carried out by flash chromatography on silica with medium-pressure liquid chromatography (MPLC; *Biotage*® Selekt System) and on C-18 by reversed-phase high-performance liquid chromatography (RP-HPLC; *Agilent* 1290 Infinity II System).

#### Methods

##### Mass spectrometry

Reaction monitoring and fractionation patterns were measured on an Atmospheric Solid Analysis Probe (ASAP) mass spectrometer (Model: *Expression Compact*, Advion). Atmospheric pressure chemical ionization (APCI) was used as ionization method.

Liquid chromatography electrospray-ionization mass-spectrometry (LC-ESI-MS) was measured on *1290 Infinity II* ultra-high performance liquid chromatography (UPLC, *Agilent,* USA) connected to a single quadrupole mass spectrometer (SQ/MS; module type: G6125C; *Agilent,* USA).

High-resolution mass spectrometry (HRMS) was measured by the staff of the Mass Spectrometry Laboratory of the Department of Chemistry at the University of Zurich employing a Dionex Ultimate 3000 UHPLC system (*Thermo-Fischer Scientific*) connected to a QExactive MS with a heated ESI source (*Thermo-Fisher Scientific*).

##### NMR spectroscopy

NMR data were recorded on an AV2-400 or AV2-500 NMR spectrometer (Bruker). The samples were measured in Norell® natural quartz (5 mm, 500 MHz) NMR tubes at ambient temperature. The chemical shift is given in ppm relative to the internal standard tetramethylsilane ($^1H$: $\delta(SiMe_4) = 0.00$) or relative to the solvent reference signal ($^1H$: $\delta(CDCl_3) = 7.26$ ppm or $\delta(CD_3CN) = 1.94$ ppm;

$^{13}C$: $\delta(CDCl_3)$ = 77.16 ppm or $\delta(CD_3CN)$ = 1.32 and 118.26 ppm). The multiplicities were given as follows: singlet (s), doublet (d), triplet (t), quartet (q), multiplet (m), centered multiplet ($m_c$) and broad singlet (brs).

The mechanism study was performed with CouCyCN **1b** (6.8-7.3 mM) in $CD_3CN$. After adding 3.3 M NaOD (40% in $D_2O$), the mixture was thoroughly mixed and incubated for 10 min at 21 °C, then the $^1H$-NMR spectra (400 MHz, 128 scans) were measured.

**Optical spectroscopic methods**

Stock solutions of chemical probes were prepared in $(CH_3)_2SO$ (DMSO spectrophotometric grade > 99.9%, *Acros Organics*) at a concentration of 20 mM and 1 mM and stored at –20 °C. All stock solutions were thawed immediately before use. UV-visible spectra were measured on a Multiskan SkyHigh microplate spectrophotometer (*Thermo-Fisher* Scientific) using *SkanIt* software. Fluorescence spectra were acquired using an FS5 spectrofluorometer (*Edinburgh* Instruments) equipped with an SC-25 cuvette holder or SC-40 plate reader using *Fluoracle* software. The obtained spectra were background corrected.

Absolute fluorescence quantum yields were measured at concentrations of 1 µM by employing an SC-30 integrating sphere. Extinction coefficients were determined in a dilution series of 2.5-15 µM in PBS (1x, pH 7.4), measuring the absorbance at $\lambda_{max}$ and applying a linear regression. All measurements were conducted at ambient temperature and under red light ambient illumination with 96-well plates (*Corning*) or quartz cuvettes (*Strana Scientific Ltd*, 10 mm path length). If not stated otherwise, measurements were carried out as preparative triplicates. The data analysis was performed using *Prism10 GraphPad*.

**pH profile**

The instrument pHenomenal 1100L (*VWR*) was calibrated before every use. pH buffers were adjusted by dropwise addition of 0.1-1 M HCl or 0.1-1 M NaOH under stirring at room temperature.

The pH profiles were determined with a 10 µM probe (1 mM in DMSO) in buffers of pH 2–13 (pH 2, citric acid; pH 3–7, citric acid/$Na_2HPO_4$; pH 8, $Na_2HPO_4$; pH 9–11, $NaHCO_3$/$Na_2CO_3$; pH 12–13, KCl/NaOH; or pH 5.5-8.5, 10x PBS). The mixture was equilibrated for 30–60 min at 37 °C. Absorbance spectra were measured in preparative triplicates, including a blank measurement without a probe. Measurements were background corrected and averaged.

**Selectivity assay**

The selectivity assays were performed in a total volume of 200 µL with 10 µM CouCyCF$_3$ **1b** or CouCyCN **1c** (1 mM in DMSO) at pH 5.5 and 7.0 in 10x PBS. The following analytes in biological relevant concentrations were used: 1 µM $HS^{-}$ (nM-µM range[1]), 1 mM reduced glutathione (GSH, about 1 mM in Gram-negative, but rarely in Gram-positive bacteria[1]), 200 µM $H_2O_2$ (4000-fold excess, 50 nM steady-state concentration in *E. coli*[2]), 200 µM of the nucleophilic amino acids (AA) L-cysteine, L-serine, L-lysine, L-histidine (concentrations in nM-mM range depending on AA, growth conditions and bacterial species[3]), 10 mM $Mg^{2+}$ (mM range[4,5]), 10 µM $Cu^{2+}$ (µM range[4]), 200 µM $Zn^{2+}$ (concentration required for *E. coli* growth[6]), 10 µM $Fe^{2+}$ and $Fe^{3+}$ (µM range[6]) and 10 µM $CN^{-}$ (toxic for most bacteria; cyanogenic bacteria like *Pseudomonas aeruginosa* can produce $CN^{-}$ in µM range[7]). As a blank, the analytes were measured at pH 5.5 and 7.0 in 10x PBS. As positive controls, the absorbance of CouCyCF$_3$ **1b** or CouCyCN **1c** was measured at pH 5.5, 7.0, and 8.0. The 96-well plate was incubated at 37 °C and 180 rpm for 30 min. Absorbance spectra were measured, background corrected, and triplicates averaged. The ratio $A_{415}/A_{600}$ or $A_{415}/A_{585}$ for CouCyCF$_3$ **1b** or CouCyCN **1c** was calculated, respectively.

**Thiol competition assay**

The assay was performed in a 96-well plate with a total volume of 200 µL with 5 µM CouCyCF$_3$ **1b** or CouCyCN **1c** (1 mM in DMSO) at pH 5.0, 6.0, 7.0, and 8.0 (citric acid/$Na_2HPO_4$ buffers). Stock solutions of NaSH (100 mM) and GSH (250 mM), as well as a dilution series in water, were freshly prepared. At each pH, the thiols NaSH (0.00, 0.01, 0.10, 0.50 mM) and GSH (0.00, 0.01, 0.10, 0.50, 1.00, 5.00) were added at different concentrations and incubated at 22 °C and 180 rpm for 30 min. Absorbance spectra were measured as preparative triplicates, background corrected, and averaged. The ratio $A_{415}/A_{600}$ or $A_{415}/A_{585}$ for CouCyCF$_3$ **1b** or CouCyCN **1c** was calculated, respectively.

**Kinetics measurements**

Kinetic studies were performed using solutions of CouCyCF$_3$ **1b** and CouCyCN **1c** (5 µM in ddH$_2$O, pH 5.8) under pseudo-first-order conditions. The absorbance decrease or increase at $\lambda_{max}$ was measured after the addition of an excess of NaOH or HCl, respectively. A solution of the chemical probes in ddH$_2$O (pH 5.8) was prepared in a quartz cuvette and placed in a spectrophotometer. The measurement was started and the absorbance at $\lambda_{max}$ was measured for 20 seconds, then 10 equiv. NaOH (100 µL of aqueous 1 mM NaOH solution) was added and quickly mixed. The emission at $\lambda_{max}$ was measured every 10 seconds over 3 minutes. A new measurement was started, $\lambda_{max}$ was measured for 20 seconds and 300 equiv. HCl (100 µL of aqueous 30 mM HCl solution) was added to the same cuvette to observe the reversible reaction. The absorbance at $\lambda_{max}$ was measured every 10 seconds over 4 minutes. All measurements were performed as preparative triplicates at ambient temperature. Data were averaged to obtain a mean with standard deviation and fit to an exponential plateau function to calculate the kinetic constant $k_{obs}$ ($s^{-1}$) and the goodness of the fit $R^2$.

**Polarity profile**

The polarity-dependent emission spectra were measured in mixtures containing 0% ($\varepsilon$ = 80.38), 10% ($\varepsilon$ = 72.02), 20% ($\varepsilon$ = 63.50), 40% ($\varepsilon$ = 45.96), 60% ($\varepsilon$ = 28.09), 80% ($\varepsilon$ = 12.19), or 100% ($\varepsilon$ = 2.24) volume-% dioxane in water.[8] The emission spectra of the probes (0.5 µM) were measured in glass quartz cuvettes at the blue and red emission maxima with a spectrofluorometer. The spectra were measured as preparative triplicates, background-subtracted, and averaged. The emission maxima of each mixture were plotted against the dielectric constant and interpolated as a Gaussian curve.

**Laboratory bacterial strains and standard cultivation conditions**

*E. coli* K12 was obtained from the laboratory of Prof. Alexandre Persat (EPFL), and *S. epidermidis* WDCM 0036 Vitroids™ was purchased from *Sigma-Aldrich*. *E. coli* BW25113 parental strain and the Δ*cfaS* knockout strain carrying a *cfa::kanamycin* mutation (clone ID: JW1653) were supplied from the Keio Knockout Collection.[9,10]

Cultivation of bacteria without a resistance gene was conducted at 37 °C and 180 rpm for 16 h in culture tubes in 5 mL lysogeny broth (LB) medium, or on LB agar plates (LB medium completed with 15 g $L^{-1}$ agar). The Δ*cfaS E. coli* strain was cultured in LB containing 50 mg $L^{-1}$ kanamycin (kan). The plasmid pSCM001 (Addgene: 124605)[11] expresses the mCherry-pHluorin fusion protein under an arabinose-inducible promoter and was transformed into chemically competent *E. coli* K12 cells. Bacteria were cultured at 37 °C, at 180 rpm in LB medium containing 100 µg $mL^{-1}$ carbenicillin to maintain the plasmid, and 5 mg $L^{-1}$ arabinose to induce protein expression. Cell growth and viability were assessed with optical density (OD) measurement and colony-forming units (CFU) on agar plates.

**Cultivation of clinical isolates**

The collection, pseudonymization, and usage of clinical samples used in this project was carried out with the appropriate ethics approval within the NCCR AntiResist project (2020-02588). Methicillin-resistant *S. aureus* (MRSA, t619915) strain was isolated from a prosthetic joint infection, and a persistent methicillin-sensitive *S. aureus* (MSSA, P70) strain from a deep-seated infection and bacteremia at the University Hospital of Basel, Switzerland. The *S. aureus* strains were incubated in RPMI 1640 (*Thermo Fisher Scientific*) supplemented with 10% fetal bovine serum (FBS, *Thermo Fisher Scientific*) at 37 °C when grown in liquid culture or on Columbia Blood Agar plates (*Becton Dickinson*, p/n 254071).

**THP-1 cultivation conditions**

THP-1 monocytes (passage 2-8) were cultured in T25 (10 mL culture) or T75 (20 mL culture) flasks at 37 °C and 5% $CO_2$. As culture medium RPMI 1640 (*Thermo Fisher Scientific*) without phenol red, supplemented with 10% fetal bovine serum (FBS, *Thermo Fisher Scientific*) was freshly prepared. Cells were subcultured every two to three days at a concentration of $9 \times 10^5$ to $1 \times 10^6$ cells $mL^{-1}$. Fresh cultures were seeded by transferring the cell suspension into pre-warmed culture medium to seed 4-5 × $10^5$ cells in a total volume of 20 mL (T75). Cell viability was assessed using trypan blue (*Thermo Fisher Scientific*), and viable cells were counted using a hemocytometer TC20 (*Bio-Rad*).

**Blood cell cultivation conditions**

Approximately 7 mL of fresh heparinized whole blood was collected from healthy donors at the University Hospital of Basel, Switzerland. Blood was washed with phosphate-buffered saline (PBS), and the resulting cell pellet was resuspended in 1 mL of red blood cell lysis buffer (*Thermo Fisher Scientific*, p/n 00-433-57) for 5 minutes at room temperature. Following lysis, 50 mL of PBS was added, and cells were centrifuged at 350 × g for 5 minutes. The pellet was resuspended in 1 mL of RPMI 1640 medium supplemented with 10% FBS, and viable cells were counted using a TC20 automated cell counter (*Bio-Rad*). Cells were seeded at a density of $2 \times 10^5$ cells per well in flat-bottom 96-well plates and incubated under standard conditions (37 °C, 5% $CO_2$) prior to infection.

**Preparation of poly-*D*-lysine coated glass chamber slides for imaging**

8-well plate chambered cover glass slides (*ibidi*, Germany) were coated with 250 µL of poly-*D*-lysine (50 µg $mL^{-1}$ in $ddH_2O$) and incubated at room temperature for 30 min. The poly-*D*-lysine solution was aspirated, and the chamber slides were washed three times with 400 µL $ddH_2O$ to remove excess poly-*D*-lysine. The excess $ddH_2O$ was aspirated after the final wash and the slides were allowed to air-dry at room temperature under UV light for 1–1.5 h.

**Bacterial imaging protocol**

A single bacterial colony was incubated in 5 mL LB at 37 °C, shaking at 180 rpm for 16 h. Using the overnight culture, a fresh culture was inoculated with an $OD_{600}$ of 0.1 in LB and incubated at 37 °C and 180 rpm until an $OD_{600}$ of 0.6–0.7 was reached (approx. 2 h). Two aliquots of 200 µL were prepared in fresh microcentrifuge tubes. The cells were centrifuged at 10’000 rpm for 1 min at room temperature. The LB media was removed, and the pellet was suspended in 1 mL imaging medium. The cells were pelleted at 10’000 rpm for 1 min at room temperature, the supernatant was carefully removed, and the pellet was resuspended in 1 mL imaging solution. To each well on a poly-*D*-lysine coated 8-well chamber slides, 200 µL cell suspension was added and incubated at 37 °C for 20 min to promote the adhesion of bacterial cells to the surface. The chambers were gently washed three times with 400 µL imaging medium to remove unattached bacteria cells. For imaging with chemical probes,

200 µL imaging medium containing 1–5 µM probe (1 mM stock solution in DMSO) was added and incubated for 10 min at 37 °C.
For pH calibration experiments with chemical probes and pHluorin, the solution was aspirated, and 250 µM carbonyl cyanide *m*-chlorophenyl hydrazone (CCCP; 50 mM stock solution in DMSO; *Acros Organics or Sigma-Aldrich*) in PBS (10x) with defined pH was added. Before imaging, the bacterial cells were incubated for 30–60 min to equilibrate external and internal pH.

FluoroBrite™ DMEM (high *D*-Glucose, 3.7 g $L^{-1}$ sodium bicarbonate; pH 7.4; *Thermo-Fisher Scientific*) was used as imaging medium. For the pH calibration experiment, PBS (10x) was manually adjusted to pH 5.0, 5.5, 6.0, 6.5, 7.0, 7.5, 8.0, and 8.5 and filtered with a syringe filter (0.22 µm). For experiments with pHluorin, all bacterial cultures in LB were supplemented with 100 µg $mL^{-1}$ carbenicillin and 5 mg $L^{-1}$ arabinose.

**THP-1 and blood cell coculture imaging protocol**

THP-1 monocytes (passage 2-8) or blood cells were plated on a poly-*D*-lysine-coated 8-well chamber slide at a density of 50’000 cells in 250 µL per well. The cells were allowed to settle for 15 min before the coculture was initiated.
$1.0 \times 10^7$ cells of an *S. aureus* or *S. epidermidis* overnight culture were transferred to a 1.5 mL tube and centrifuged at 10’000 rpm for 1 min. The media was removed, the pellet resuspended in 1 mL imaging solution, and pelleted at 10’000 rpm for 1 min. This step was repeated twice. The final pellet was resuspended in 1 mL imaging solution containing 1 µM CouCyCF$_3$ **1b**. After incubation at 37 °C for 10 min protected from light, the cells were pelleted (10’000 rpm, 1 min) and washed with imaging solution (3 × 1 mL). The final bacterial suspension was further diluted to $2.0 \times 10^5$, $6.0 \times 10^5$, and $1.2 \times 10^6$ cells $mL^{-1}$. To induce the coculture, the medium from the plated THP-1 or blood cells was gently removed, and 250 µL of the bacterial dilution was added to obtain the desired multiplicity of infection (MOI) of 1, 3, or 6. FluoroBrite™ DMEM (pH 7.4) or 1x PBS (pH 7.4) was used as imaging medium as indicated in each experiment.

**Confocal microscopy**

Imaging was performed with a *Nikon* W1 spinning disk microscope equipped with a dual-camera system (CMOS, Photometrix). Fluorescence images were collected using a CFI Plan Apochromat Lambda D oil-immersion objective (60x, NA = 1.4). Brightfield images were taken with a white light-emitting diode, while fluorescence images were performed with laser lines and filters as appropriate. Channels were imaged sequentially. The microscope was operated using *Nikon* NIS-elements software. Imaging was performed at 37 °C.

**Image analysis**

Imaging processing and analysis were performed with *Fiji – Image J* software. Ratiometric images were obtained by dividing the two channels of interest (CouCy's: $I_{640}/I_{445}$; mCherry-pHluorin: $I_{488}/I_{561}$; BCECF-AM: $I_{488}/I_{455}$). A mask of each field of view (FOV) was created using a threshold image (Li method). The obtained binary image was further processed if required (e. g., to separate single cells if possible). Single bacteria cells and clustered bacteria cells were analyzed as particles (>1 $\mu m^2$; circularity 0–1) to define regions of interest (ROI). The ROI mask was applied to the ratiometric image using the ROI manager. The mean of all ratiometric pixels per bacterial cell or cluster ROI was calculated and represents one data point. For automation, the described processing was performed using a custom-written macro. Further statistical analysis was performed using *Prism10 GraphPad*. The median and standard deviation were calculated for the linear calibration, and a linear regression was performed.

**Flow cytometry analyzer**

Flow cytometry was performed with a *BD FACSymphony 5L* equipped with five lasers (355, 405, 488, 561, and 640 nm) controlled via the *BD Coherent Connect* software. 100,000 events per sample were recorded with the following setup: FSC, 500 V, threshold: 200; SSC, 229 V, threshold: 200; ex. 405 nm, em. 515/20 nm, 600 V; ex. 639 nm, em. 670/30 nm, 650 V. Prior to the measurement, the samples were diluted in the corresponding buffer and filtered through a cell strainer cap (mesh size 30-50 µm) by pipetting into a fresh *Falcon*® FACS tube (p/n 352235) to obtain a 7,000 to 10,000 event-per-second (evts $s^{-1}$) threshold rate and an electronic abort rate not higher than 250-300 evts $s^{-1}$. A buffer sample and an unstained *E. coli* culture were used to identify the gates for buffer salt crystals and bacteria in the logarithmic scale SSC-H vs. FSC-H plot. The bacterial population was gated for single cells first in the FSC-H vs. FSC-A and subsequently in the SSC-H vs. SSC-A plot. The emission intensities at 670/30 nm and 515/20 nm were analyzed as histograms, scatterplots, or contour plots (5-10% event density). The instrument was operated using *FACSDiva* software, and data analysis was performed using *FlowJo*.

**pH calibration experiment**

A single colony *E. coli* K12 overnight culture was used to inoculate a fresh culture in LB and was incubated at 37 °C and 200 rpm until an $OD_{600}$ of 0.8-1.0 was reached. The culture was transferred in a fresh microcentrifuge tube (8x1.0 mL) and centrifuged at 10’000 rpm for 1 min at room temperature. The LB media was removed, and the pellet was suspended 10x PBS (pH 5.0, 5.5, 6.0, 6.5, 7.0, 7.5, or 8.0) containing 250 µM CCCP (50 mM in DMSO) and 1 µM $CouCyCF_3$ **1b** or CouCyCN **1c** (1 mM in DMSO) in a total volume of 1 mL. An unstained sample was prepared as a negative control by suspending a cell pellet in 10x PBS (pH 7.4). Each sample was filtered through a cell strainer cap (mesh size 30-50 µm). The tubes were protected from light, incubated at 180 rpm and 37 °C for 60 min, and then placed on ice until the measurement. The samples were diluted at 1:20 or 1:30 with the corresponding 10x PBS buffer, and the measurement was performed as described above.

**pH-sensitive knockout strains**

A single colony of *E. coli* BW25113 (parent strain) and Δ*cfaS* knockout strain was cultured in LB (+kan for Δ*cfaS*) at 37 °C and 200 rpm overnight. The overnight culture was used to inoculate a fresh culture in LB (+kan for Δ*cfaS*) and was incubated at 37 °C and 200 rpm until an $OD_{600}$ of 0.6-0.8 was reached. The cultures were transferred to fresh 15 mL Falcon tubes and centrifuged at 10’000 rpm for 1 min at room temperature. The LB media was removed, and the pellet was suspended in 60-100 µL LB medium (pH 7.0). The samples were stored on ice until the measurement. Right before the measurement, 10 µL of the cell suspension was diluted with 1 mL LB with the corresponding pH (3.0, 4.0, 5.0, or 7.0). The cells were stained with 1 µM $CouCyCF_3$ **1b** (1 µL of 1 mM in DMSO). For the 0 min incubation measurement,

100 µL of the culture was immediately diluted (1:10) with 900 µL 1x PBS (pH 3.0, 4.0, 5.0, or 7.0), filtered, and measured as described above. The samples diluted in 1x PBS were discarded after the measurement. The cultures subjected to acid shock in LB (pH 3.0, 4.0, 5.0, or 7.0) were incubated for 10, 20, and 30 min at room temperature, protected from light. At each time point, 100 µL of the sample was filtered, diluted in the corresponding 1x PBS buffer, and measured immediately.
The pH-adjusted LB medium (pH 3.0, 4.0, 5.0, and 7.0) and 1x PBS buffer (3.0, 4.0, 5.0, and 7.0) were prepared freshly.

**THP-1 and blood cell coculture with ImageStream**

Imaging-based flow cytometry was performed using the *Amnis ImageStream*® X Mk II system equipped with three objectives (20x, 40x, and 60x), 12 detection channels, and four excitation lasers (405, 488, 561, 642 nm). Data acquisition and analysis were conducted using *IDEA*® (Image Data Exploration and Analysis Software), with gating strategies applied to single, in-focus cells.
THP-1 monocytes (passage 2-8) or blood cells were seeded at a density of $2 \times 10^5$ cells per well (*Falcon*® 96-well clear flat bottom TC treated culture microplate) in 200 µL medium.
1 mL of an overnight culture of *S. aureus* in RPMI 1640 (*Thermo Fisher Scientific*) supplemented with 10% FBS (*Thermo Fisher Scientific*) was transferred to a 1.5 mL tube and centrifuged at 5’000 × g for 5 min. The bacterial pellet was washed twice with 1x PBS (pH 7.4). The final pellet was resuspended in 1 mL 1x PBS containing 1 µM CouCy$CF_3$ **1b**. After incubation at 37 °C for 15 min protected from light, the cells were pelleted (5’000 × g, 5 min) and washed twice with 1x PBS. The final bacterial pellet was suspended in culture medium and added to the THP-1 or blood cells to obtain the desired MOI. As a reference sample, the stained *S. aureus* monoculture was analyzed. Right before the measurement, the cell coculture was transferred from the well to a 1.5 mL tube and loaded in the *ImageStream* for data acquisition.

**Toxicity assay**

The toxicity studies were performed with MRSA and MSSA with varying concentrations of CouCy$CF_3$ **1b** and CouCyCN **1c**. Bacterial strains were cultured overnight at 37 °C on Columbia Blood Agar plates (*Becton Dickinson*, p/n 254071). Single colonies were picked and resuspended in sterile medium to a McFarland standard of 0.5 ($OD_{600}$ = 0.063), then further diluted to a final concentration of approx. $2 \times 10^6$ CFU $mL^{-1}$ in 96-well plates. For $OD_{600}$ measurements, freshly diluted bacterial suspensions were plated in quadruplicate (technical replicates) in a flat-bottom 96-well microplate, with varying concentrations of CouCyCF$_3$ **1b** and CouCyCN **1c**. Peripheral wells were filled with sterile PBS to minimize edge effects due to evaporation. Plates were incubated at 37 °C for 24 hours in a BioTek Synergy™ H1 microplate reader. The plate was shaken for 3 seconds every 10 minutes prior to $OD_{600}$ readings. To assess bacterial viability, CFU per mL was determined at 0 h, 1 h, 2 h, and 24 h by serial dilution and plating on agar plates as two biological replicates.

#### Synthesis procedures

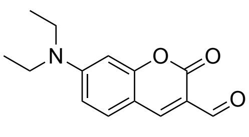
**3**
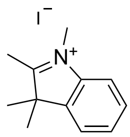
**2a**
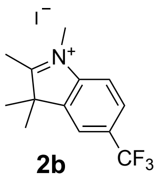
**2b**
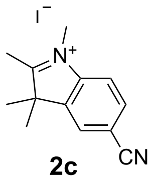
**2c**

Coumarin-aldehyde **3**[12], CouCyH **1a**[13], and indoleninium salts **2b,c**[14–17] were prepared according to published procedures. Compound **2a** is commercially available.

#### General procedure to prepare CouCy's

Aldehyde **3** (1.0 equiv.) and indoleninium salt (1.2 equiv.) were dissolved in anhydrous ethanol (0.1 mmol/mL) and the mixture was heated to 65 °C. After a few minutes, the yellow mixture turned green and finally dark blue. The reaction progress was monitored by LC-MS. After complete conversion of aldehyde **3**, the mixture was allowed to reach room temperature, and the solvent was evaporated. The residue was dissolved in $CHCl_3$, dry-loaded on Celite, and purified by flash column chromatography (12 g silica, 0–10% methanol in $CH_2Cl_2$), followed by reverse-phase column chromatography using HPLC (10–90% acetonitrile in $H_2O$ + 0.1% TFA). CouCy dyes were obtained as dark blue solids.

**(*E*)-2-(2-(7-(diethylamino)-2-oxo-2*H*-chromen-3-yl)vinyl)-1,3,3-trimethyl-5-(trifluoromethyl)-3*H*-indol-1-ium (1b)**

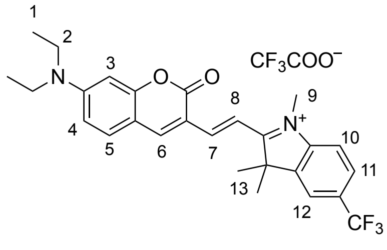

Chemical Formula: $C_{27}H_{28}F_3N_2O_2^+$
Molecular Weight: 469.5

CouCyCF$_3$ **1b** was synthesized according to the general procedure to yield a dark blue solid (116 mg, 194 µmol, 96%).

**$^1$H-NMR** (400 MHz, $CD_3CN$): $\delta$ = 1.24 (t, $^3J_{1,2}$ = 7.2 Hz, 6H, $CH_3$-1), 1.80 (s, 6H, $CH_3$-13), 3.57 (q, $^3J_{2,1}$ = 7.1 Hz, 4H, $CH_2$-2), 3.89 (s, 3H, $CH_3$-9), 6.60 (d, $^{meta,4}J_{3,4}$ = 2.4 Hz, 1H, H-3), 6.85 (dd, $^{ortho,3}J_{4,5}$ = 9.2 Hz, $^{meta,4}J_{4,3}$ = 2.5 Hz, 1H, H-4), 7.53 (d, $^{ortho,3}J_{5,4}$ = 9.2 Hz, 1H, H-5), 7.73 (d, $^{ortho,3}J_{10,11}$ = 8.4 Hz, 1H, H-10), 7.85 (d, $^{trans,3}J_{7,8}$ = 15.6 Hz, 1H, H-7), 7.90 (dd, $^{ortho,3}J_{11,10}$ = 8.5 Hz, $^{meta,4}J_{11,12}$ = 1.8 Hz, 1H, H-11), 8.01 (d, $^{meta,4}J_{12,11}$ = 1.5 Hz, 1H, H-12), 8.23 (d, $^{trans,3}J_{8,7}$ = 15.6 Hz, 1H, H-8) and 8.43 (s, 1H, H-6) ppm; **$^{13}$C-NMR** (101 MHz, $CD_3CN$): $\delta$ = 12.9, 26.7, 34.6, 46.4, 52.8, 97.7, 110.2, 111.4, 112.7, 113.6, 115.5, 121.1 (q, $J$ = 3.9 Hz), 123.8, 126.5, 127.7 (q, $J$ = 3.9 Hz), 130.4 (q, $J$ = 32.8 Hz), 133.9, 144.6, 146.0, 152.2, 153.5, 156.2, 159.5, 160.6 and 184.2 ppm; **$^{19}$F-NMR** (377 MHz, $CD_3CN$): $\delta$ = –62.34 (s, Ar-$CF_3$) and –76.46 (s, $CF_3COOH$) ppm; **LC-MS (ESI)**: $R_f$ = 2.33, m/z = 469.3 $[M+H]^+$; **HRMS (ESI)**: calculated for $[C_{27}H_{29}F_3N_2O_2]$: 469.20974, found 469.20997.

**(*E*)-5-cyano-2-(2-(7-(diethylamino)-2-oxo-2*H*-chromen-3-yl)vinyl)-1,3,3-trimethyl-3*H*-indol-1-ium (1c)**

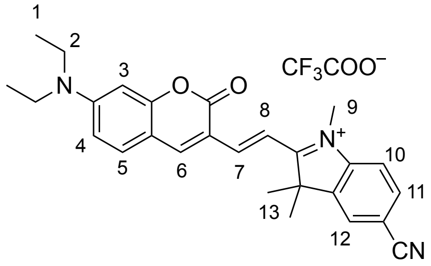

Chemical Formula: $C_{27}H_{28}N_3O_2^+$
Molecular Weight: 426.5

CouCyCN **1c** was synthesized according to the general procedure to yield a dark blue solid (103 mg, 186 µmol, 92%).

**$^1$H-NMR** (400 MHz, $CD_3CN$): $\delta$ = 1.24 (t, $^3J_{1,2}$ = 7.1 Hz, 6H, $CH_3$-1), 1.77 (s, 6H, $CH_3$-13), 3.58 (q, $^3J_{2,1}$ = 7.1 Hz, 4H, $CH_2$-2), 3.85 (s, 3H, $CH_3$-9), 6.61 (dd, $^{meta,4}J_{3,4}$ = 2.5 Hz, $^{para,5}J_{3,5}$ = 0.7 Hz, 1H, H-3), 6.87 (dd, $^{ortho,3}J_{4,5}$ = 9.2 Hz, $^{meta,4}J_{4,3}$ = 2.4 Hz, 1H, H-4), 7.54 (d, $^{ortho,3}J_{5,4}$ = 9.2 Hz, 1H, H-5), 7.69 (dd, $^{ortho,3}J_{10,11}$ = 8.5 Hz, $^{para,5}J_{10,12}$ = 0.6 Hz, 1H, H-10), 7.83 (d, $^{trans,3}J_{7,8}$ = 15.5 Hz, 1H, H-7), 7.93 (dd, $^{ortho,3}J_{11,10}$ = 8.4 Hz, $^{meta,4}J_{11,12}$ = 1.5 Hz, 1H, H-11), 8.02 (dd, $^{meta,4}J_{12,11}$ = 1.6 Hz, $^{para,5}J_{12,10}$ = 0.6 Hz, 1H, H-12), 8.24 (d, $^{trans,3}J_{8,7}$ = 15.4 Hz, 1H, H-8) and 8.45 (s, 1H, H-6) ppm; **$^{13}$C-NMR** (126 MHz, $CD_3CN$): $\delta$ = 12.9, 26.8, 34.5, 46.5, 52.6, 97.8, 109.9, 111.6, 112.0, 112.9, 113.6, 115.6, 119.2, 127.8, 134.0, 134.1, 134.8,

144.5, 146.4, 152.3, 154.0, 156.4, 159.7, 160.5 and 184.0 ppm; **$^{19}$F-NMR** (376 MHz, $CD_3CN$): $\delta$ = –75.98 (s, $CF_3COOH$) ppm; **LC-MS (ESI)**: $R_f$ = 2.13, m/z = 426.4 $[M+H]^+$; **HRMS (ESI)**: calculated for $[C_{27}H_{28}N_3O_2]$: 426.21760, found 426.21735.

#### Supplementary figures

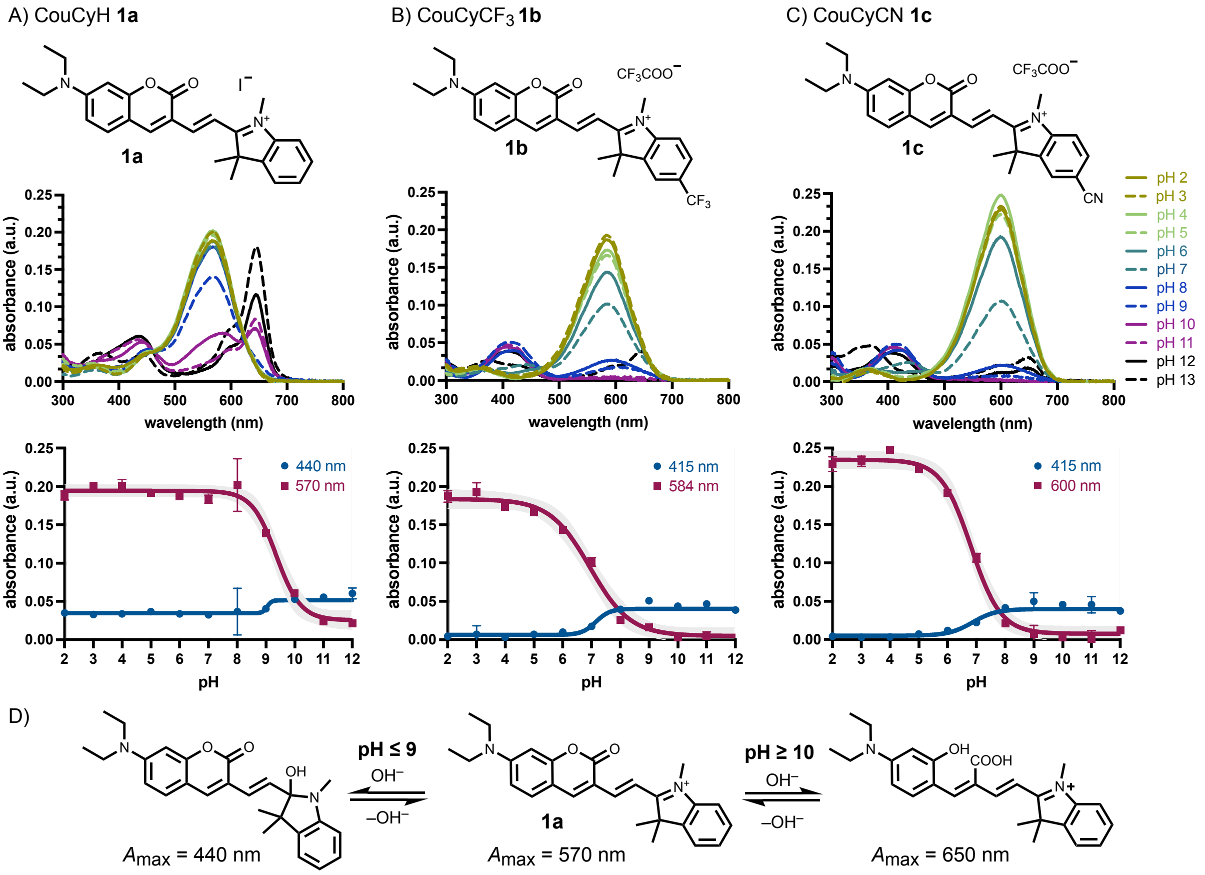

**Figure S1:** pH-dependent absorbance spectra and pH profile of A) CouCyH **1a**, B) CouCyCF$_3$ **1b**, and C) CouCyCN **1c**. Probe (5 µM) was incubated for 60 min at 37 °C. As pH buffer citric acid and $Na_2HPO_4$ (pH 2–8), $NaHCO_3$ and $Na_2CO_3$ (pH 9–11) or NaOH and KCl (pH 12–13) were used. All spectra were background-subtracted and represent the mean of preparative triplicates. Data points in the titration curve represent means and error bars are standard deviation at the indicated absorbance maxima in arbitrary units (a.u.). pH-dependent change of the red and blue absorbance maxima were interpolated as sigmoidal curves. The grey shaded area represents the confidence interval of the sigmoidal interpolation. D) Proposed mechanism of red-shifted absorbance of CouCyH **1a** with increasing pH. At high $OH^-$ concentrations (pH ≥ 10), the lactone of the coumarin core can be hydrolyzed, resulting in a longer conjugation path and red-shifted absorbance.[18] Thus, the regioselectivity of the nucleophilic attack by $OH^-$ depends on the electrophilicity of the indoleninium. Molecules with a more electron-rich indoleninium are prone to undergo this competitive nucleophilic attack on the coumarin lactone.

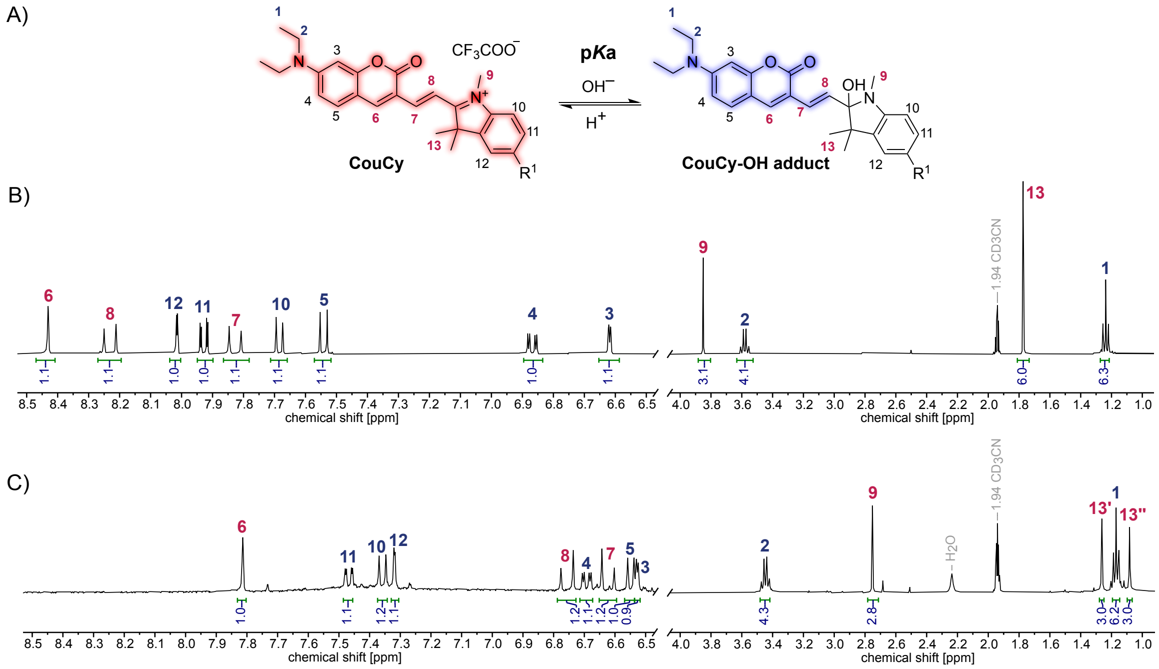

**Figure S2:** Sensing mechanism of CouCy. A) Chemical sensing equilibrium between CouCy and CouCy-OH adduct. B) $^1$H-NMR spectrum of CouCyCN **1c** (7.3 mM) in $CD_3CN$. C) $^1$H-NMR spectrum of CouCyCN **1c** (6.8 mM) in $CD_3CN$ and NaOD in $D_2O$ (3.3 M) after 10 min incubation at 21 °C.

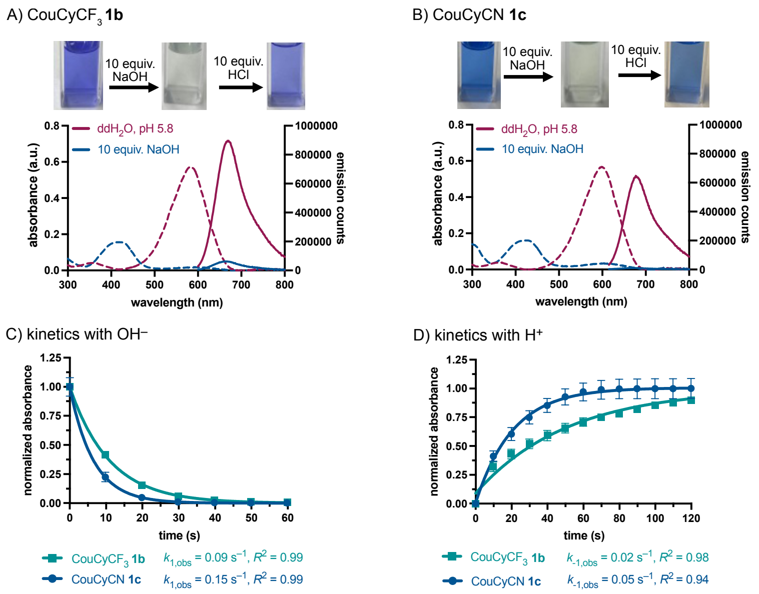

**Figure S3:** Kinetic studies with $CouCyCF_3$ **1b** and CouCyCN **1c**. A,B) Absorbance (dashed lines) and emission (solid lines) spectra of $CouCyCF_3$ **1b** (A) and CouCyCN **1c** (B) in MilliQ water (pH 5.8) before and after the addition of NaOH (10 equiv.). C) Kinetic plot of absorbance decreases after adding 100 µL NaOH (1 mM, 10 equiv.). D) Kinetic plot of absorbance increases after adding 100 µL HCl (30 mM, 300 equiv.). Absorbance was measured every 10 s, at the absorbance maxima $A_{max}$ (**1b**: 585 nm; **1c**: 600 nm). Experiments were performed as preparative triplicates, data points are means and error bars are standard deviations. The data are consistent with a pseudo-first-order reaction ($[A]=[A]_0\ e^{-k_{obs}t}$) and the kinetic constant $k_{obs}$ was calculated using a single exponential function.

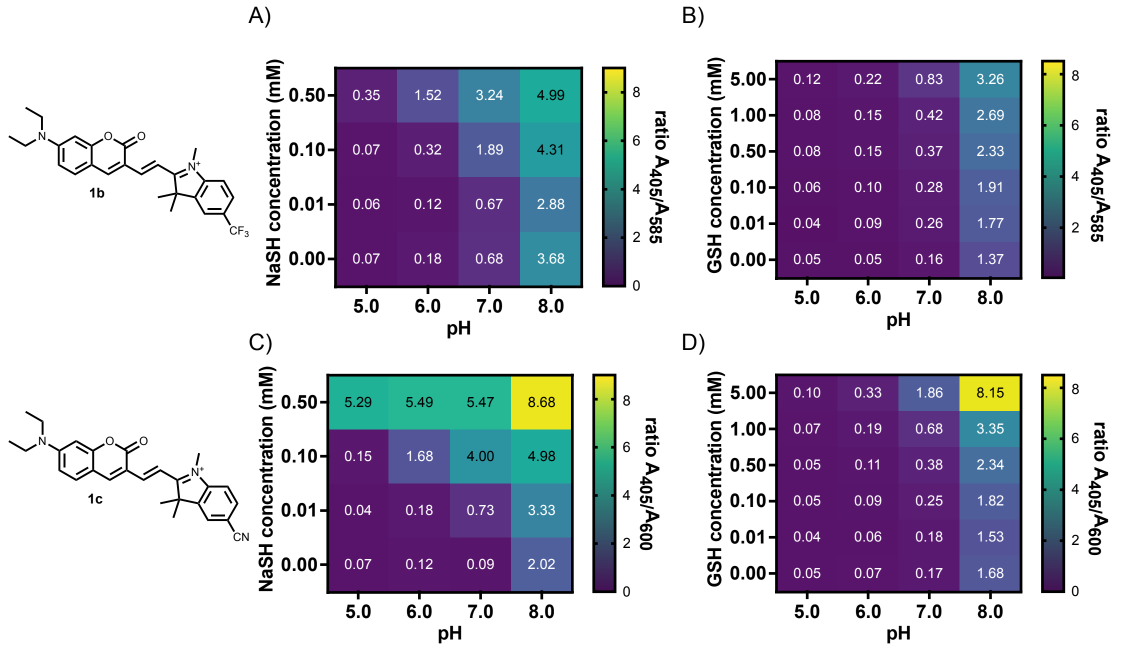

**Figure S4:** Thiol competition assay. A,B) Heatmap displaying the ratio change of CouCy$CF_3$ **1b** in an $OH^-/HS^-$ (A) and in an $OH^-$/GSH (B) sensing competition. C,D) Heatmap displaying the ratio change of CouCyCN **1b** in an $OH^-/HS^-$ (C) and in an $OH^-$/GSH (D) sensing competition. The probes (5 µM) were incubated in pH buffer (pH 5–8: citric acid and $Na_2HPO_4$) and indicated concentrations of the competing nucleophile for 30 min at 22 °C. Experiments were performed as preparative triplicates, and data were background corrected, averaged, and displayed as mean values.

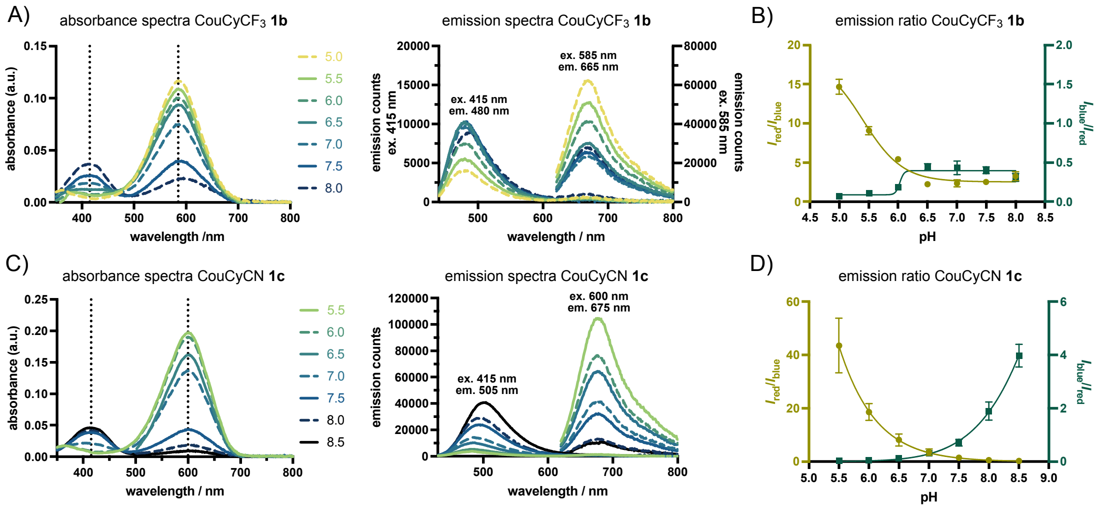

**Figure S5**: pH-dependent absorbance and emission spectra in 10x PBS. A) CouCy$CF_3$ **1b** pH-dependent absorbance spectra (left; 5 µM probe) and emission spectra (right; 1 µM probe) with two y-axis (ex. 415 nm and ex. 585 nm). B) Dynamic range of CouCy$CF_3$ **1b** in aqueous medium based on the emission ratio $I_{red}/I_{blue}$ (left y-axis) or $I_{blue}/I_{red}$ (right y-axis). C) CouCyCN **1c** pH-dependent absorbance spectra (left; 5 µM probe) emission spectra (right; 1 µM). D) Dynamic range of CouCyCN **1c** in aqueous medium based on the emission ratio $I_{red}/I_{blue}$ (left y-axis) or $I_{blue}/I_{red}$ (right y-axis). All spectra were measured in quartz cuvettes, were background-corrected, and represent the mean of preparative triplicates. The ratiometric data are plotted as means, and error bars represent standard deviations.

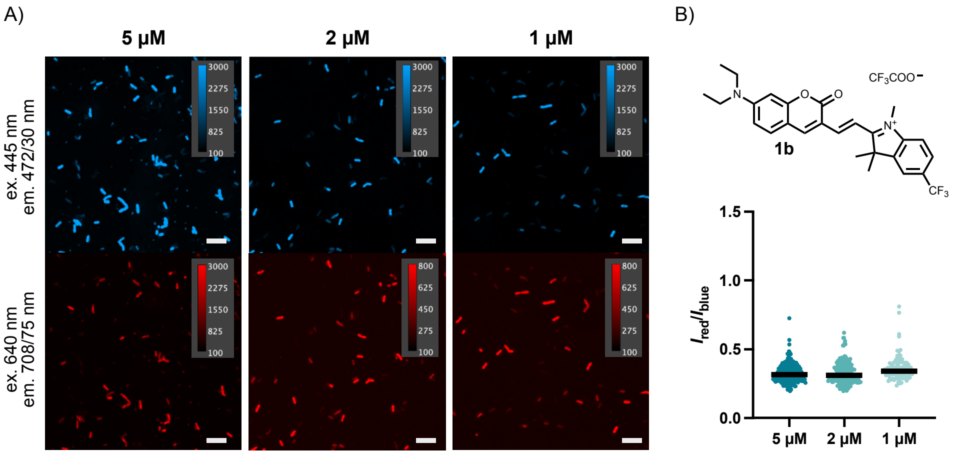

**Figure S6**: Concentration-dependent uptake of CouCyCF3 **1b** in *E. coli* K12. A) Confocal microscopy images of *E. coli* K12 stained with 5 µM, 2 µM, or 1 µM CouCyCF3 **1b** in FluoroBrite DMEM (pH 7.4). B) Scatter plot of the ratiometric analysis $I_{red}/I_{blue}$ for $N$ = 259, 183, 103 (from left to right) independent single cells or cell clusters. Imaging was performed with the following laser setup: ex. 445 nm, em. 472/30 nm, 1.3 mW, 400 ms; ex. 640 nm, em. 708/75 nm, 1.3 mW, 400 ms. Scale bar, 10 µm.

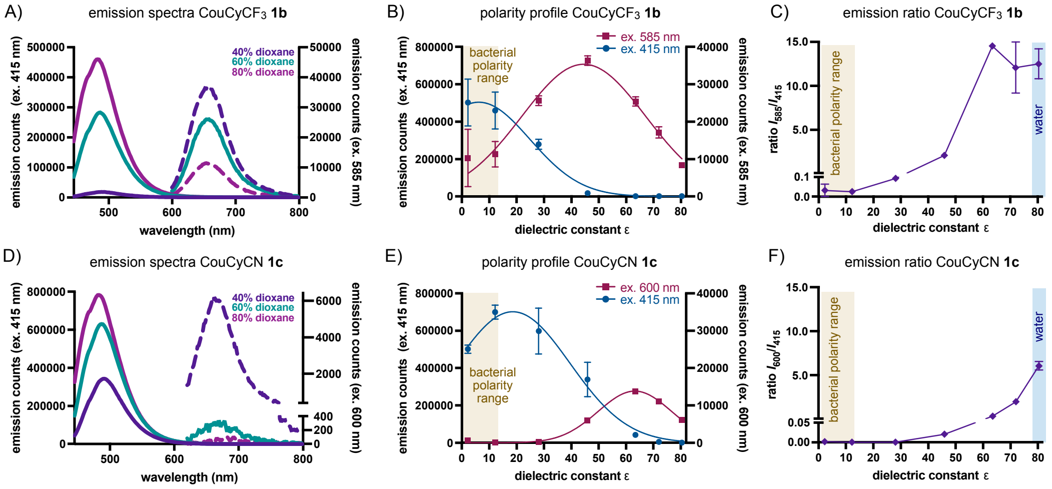

**Figure S7:** Polarity profile of CouCyCF3 **1b** and CouCyCN **1c**. A) Polarity-dependent emission spectra CouCyCF3 **1b** in 40-80% dioxane in water, excited at 415 nm (solid lines) and 585 nm (dashed lines). B) Polarity-dependent emission counts of CouCyCF3 **1b** at blue (ex. 415 nm, em. 485 nm) and red (ex. 585 nm, em. 652 nm) emission maxima. C) Polarity-dependent emission $I_{red}/I_{blue}$ ratio change of CouCyCF3 **1b**. D) Polarity-dependent emission spectra CouCyCN **1c** in 40-80% dioxane in water, excited at 415 nm (solid lines) and 600 nm (dashed lines). E) Polarity-dependent emission counts of CouCyCN **1c** at blue (ex. 415 nm, em. 495 nm) and red (ex. 600 nm, em. 664 nm) emission maxima. C) Polarity-dependent emission $I_{red}/I_{blue}$ ratio change of CouCyCN **1c**. All emission spectra of the probes (0.5 µM) were measured in quartz cuvettes, were background-corrected, and represent the mean of preparative triplicates. The emission counts at the emission maxima are plotted as mean with standard deviation against the reported dielectric constant ε of the dioxane/water mixture and interpolated as a Gaussian curve.[8] The dielectric constant ε range of bacteria is below 10 and is highlighted in beige.[19–21]

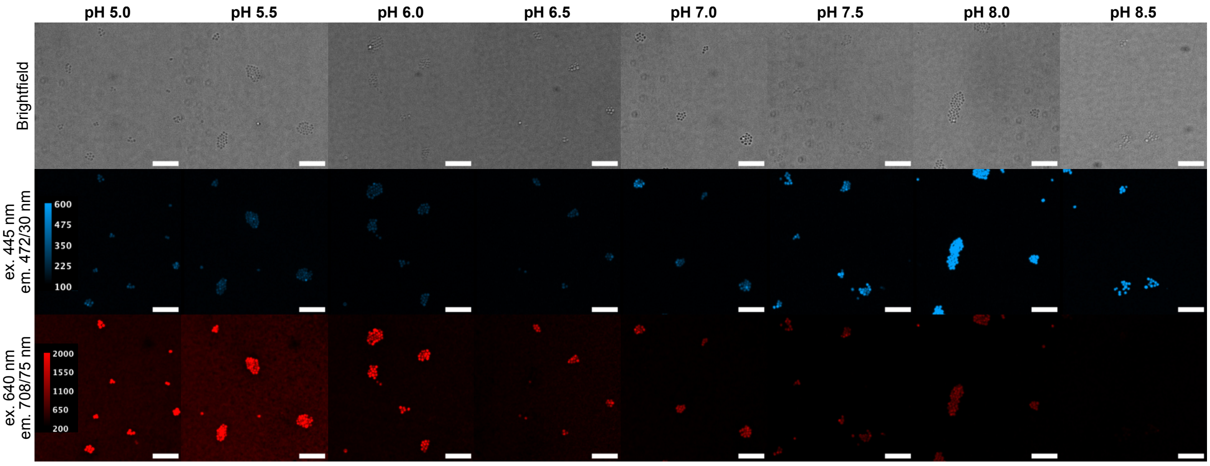

**Figure S8:** Confocal microscopy images of pH sensing in *S. epidermidis* with CouCy$CF_3$ **1b**. Bacteria were treated with CouCy$CF_3$ **1b** (1 µM) for 20 min at 37 °C, followed by a CCCP (250 µM) treatment in PBS (10x, varying pH) for 60 min at 37 °C. Imaging was performed with the following laser setup: ex. 445 nm, em. 472/30 nm, 1.8 mW, 400 ms; ex. 640 nm, em. 708/75 nm, 3.7 mW, 400 ms. Scale bar, 10 µm.

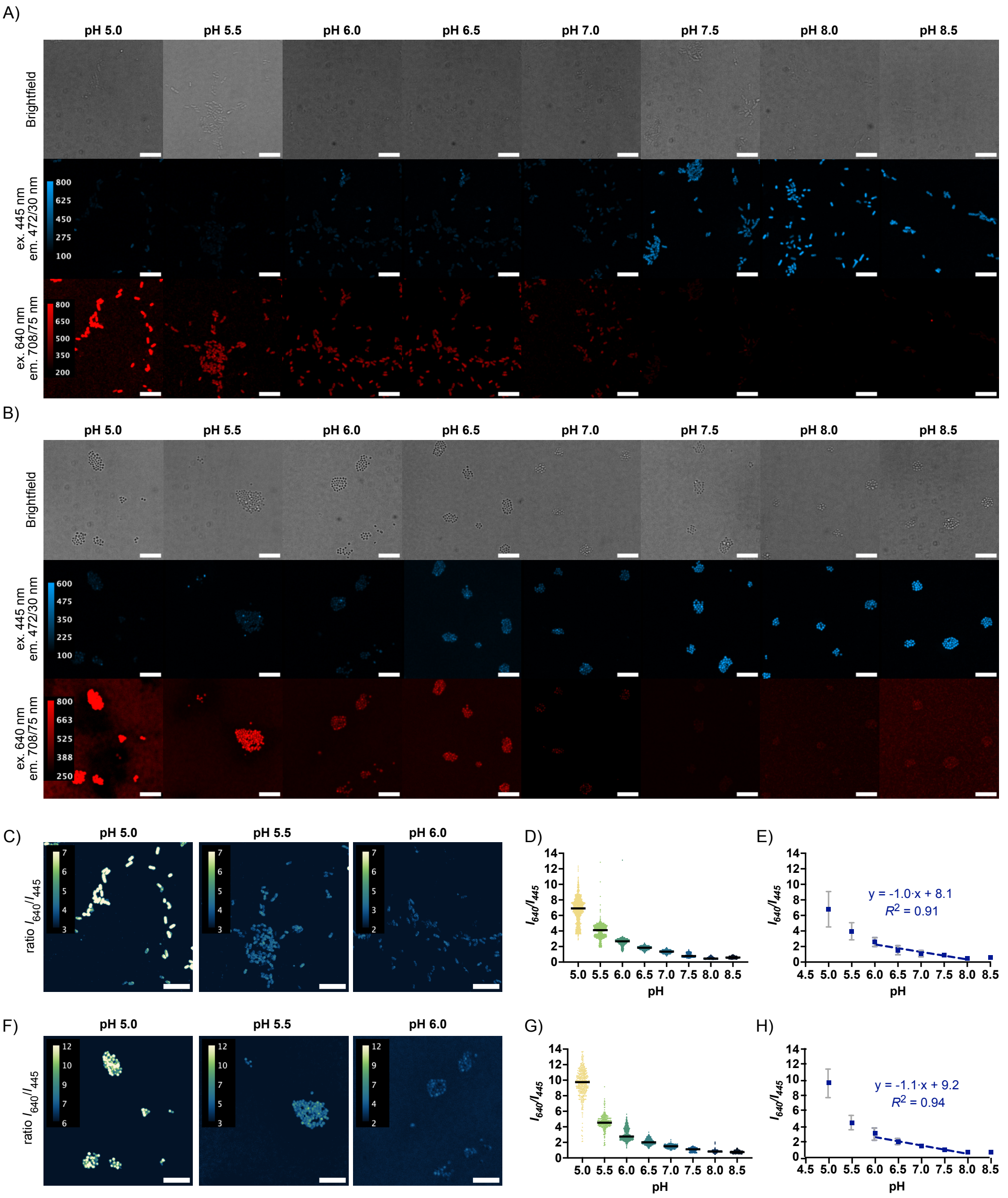

**Figure S9:** pH sensing with CouCyCN **1c**. A) Confocal microscopy images of *E. coli* K12 treated with CouCyCN **1c** (1 µM) for 20 min at 37 °C, followed by CCCP (250 µM) in PBS (10x, different pH) for 60 min at 37 °C. B) Confocal microscopy images of *S. epidermidis* treated with CouCyCN **1c** (0.5 µM) for 20 min at 37 °C, followed by CCCP (250 µM) in PBS (10x, varying pH) for 60 min at 37 °C. C) Ratiometric images of *E. coli* K12 at pH 5.0, 6.0, and 7.0. D) Scatter plot of $I_{640}/I_{445}$ ratio with $N$ = 953, 1180, 1822, 918, 803, 1014, 1104, 728 (from left to right), and E) linear regression of the $I_{640}/I_{445}$ mean with standard deviation for *E. coli* K12 imaging, of independent single cells or cell clusters from two separate imaging sessions. F) Ratiometric images of *S. epidermidis* at pH 5.0, 6.0 and 7.0. E) Scatter plot of $I_{640}/I_{445}$ ratio with $N$ = 425, 309, 577, 481, 288, 248, 1361, 211 (from left to right), and H) linear regression of the $I_{640}/I_{445}$ mean with standard deviation, for *S. epidermidis* independent single cells or cell clusters from two separate imaging sessions. Imaging was performed with the following laser setup: ex. 445 nm, em. 472/30 nm, 1.8 mW, 400 ms; ex. 640 nm, em. 708/75 nm, 3.7 mW, 400 ms. Scale bar, 10 µm.

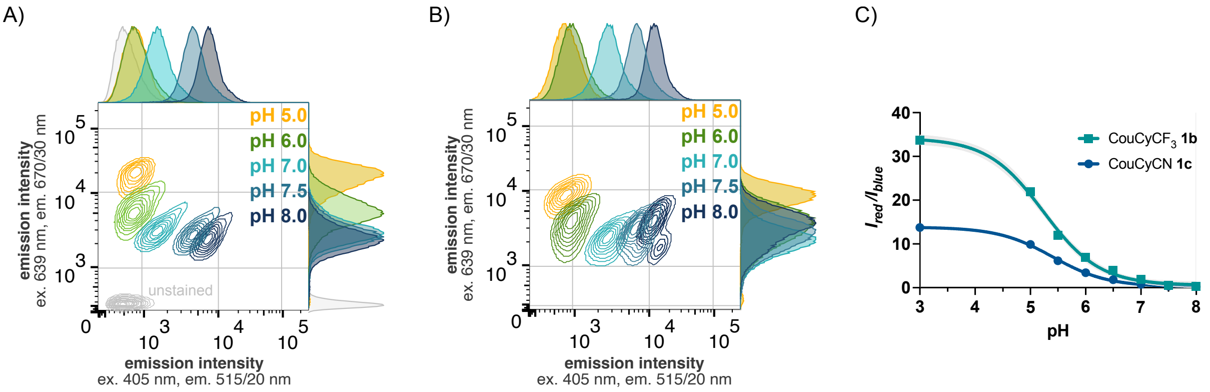

**Figure S10**: Flow cytometry pH calibration experiment. Contour plot of *E. coli* K12 cells stained with A) CouCyCF$_3$ **1b** (1 µM) and B) CouCyCN **1c** (1 µM). Each contour plot (10% event density) represents a separately measured sample and is gated for single cells (100,000 events per sample). C) Quantitative analysis of the ratio $I_{red}/I_{blue}$ against pH. The ratio was calculated with the mean intensities of each sample. Data were averaged from two independent replicates and were interpolated as sigmoidal curves. The grey shaded area represents the confidence interval of the sigmoidal interpolation.

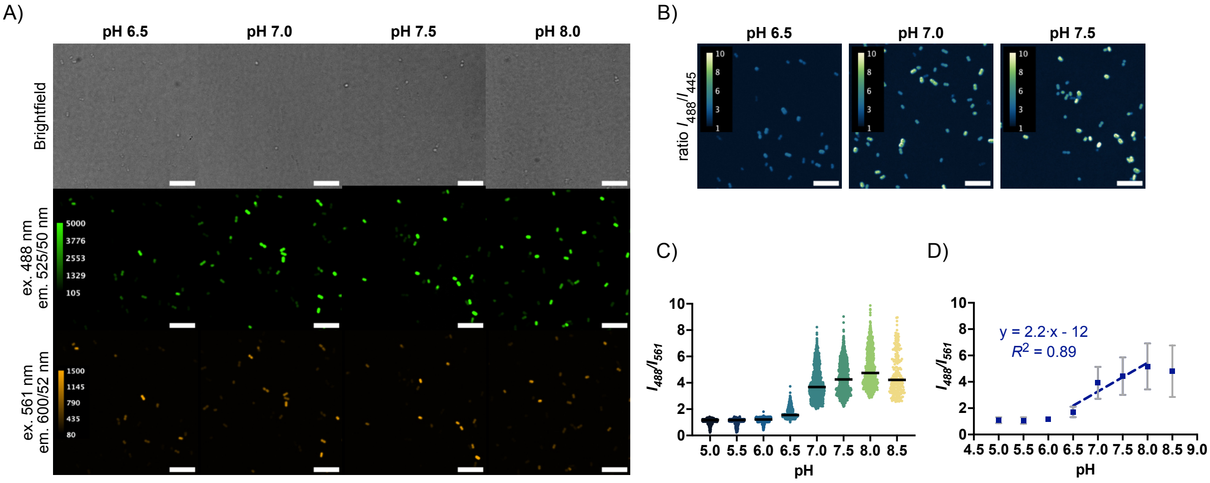

**Figure S11:** pH sensing in *E. coli* K12 expressing mCherry-pHluorin. A) Confocal microscopy images of bacteria treated with CCCP (250 µM) in PBS (10x, different pH) for 60 min at 37 °C. B) Ratiometric images at pH 6.5, 7.0, and 7.5. C) Scatter plot of $I_{488}/I_{561}$ ratio, with $N$ = 548, 584, 1321, 982, 1180, 1113, 1001, 230 (from left to right) and D) linear regression of the $I_{488}/I_{561}$ mean with standard deviation, independent single cells or cell clusters from two separate imaging sessions. Imaging was performed with the following laser setup: ex. 488 nm, em. 525/50 nm, 0.8 mW, 400 ms; ex. 561 nm, em. 600/52 nm, 0.3 mW, 400 ms. Scale bar, 10 µm.

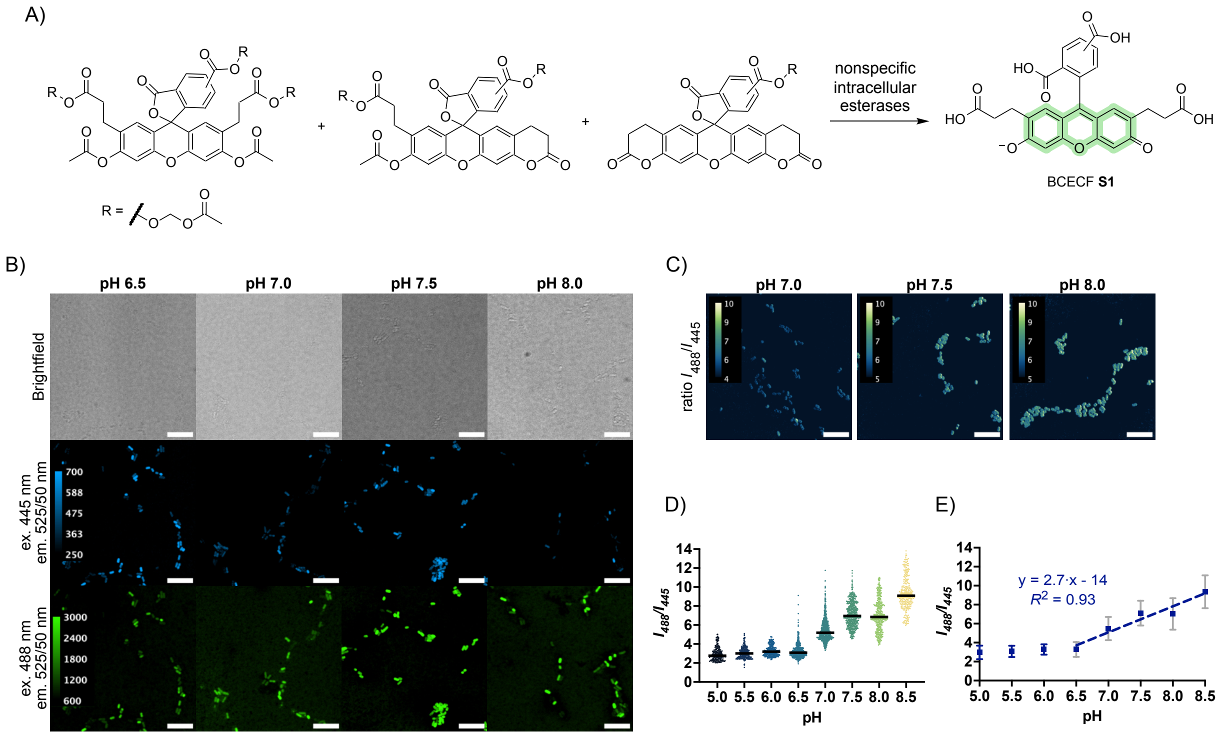

**Figure S12:** pH sensing in *E. coli* K12 with BCECF **S1**. A) Chemical structures of the acetoxymethyl (AM) ester derivatives of 2',7'-bis-(carboxyethyl)-5(6)-carboxyfluorescein (BCECF **S1**).[22] After enzymatic hydrolysis of AM esters by nonspecific intracellular esterases the pH-sensitive fluorophore BCECF **S1** is released. B) Confocal microscopy images of bacteria treated with BCECF-AM **S1** (1 µM; *Merck*, p/n 216254) for 20 min at 37 °C, followed by CCCP (250 µM) in PBS (10x, varying pH) for 60 min at 37 °C. C) Ratiometric images of *E. coli* at pH 7.0, 7.5, and 8.0. D) Scatter plot of $I_{488}/I_{445}$ ratio with $N$ = 232, 276, 316, 484, 651, 560, 560, 401, 288 (from left to right), and E) linear regression of the $I_{488}/I_{445}$ mean with standard deviation, independent single cells or cell clusters from two separate imaging sessions. Imaging was performed with the following laser setup: ex. 445 nm, em. 525/50 nm, 2.2 mW, 400 ms; ex. 488 nm, em. 525/50 nm, 2.0 mW, 400 ms. Scale bar, 10 µm.

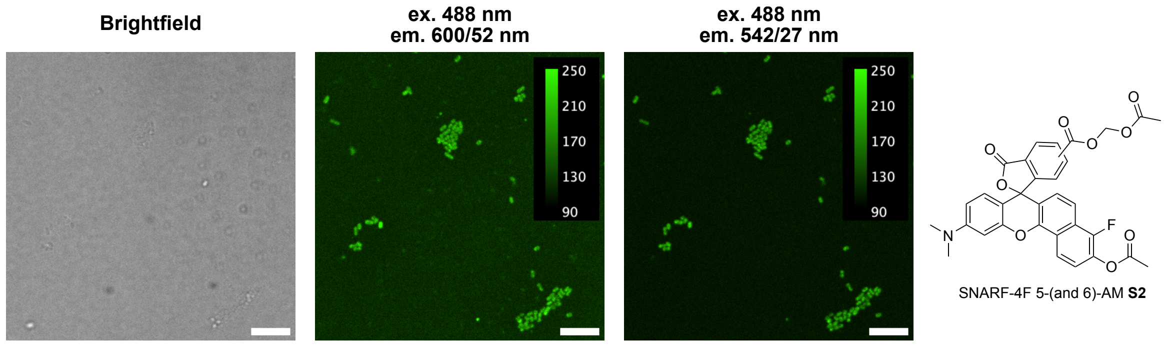

**Figure S13:** Microscopy images of *E. coli* K12 stained with SNARF-4F 5-(and 6)carboxylic acid, acetoxymethyl ester, acetate (**S2**; *Invitrogen*™, p/n S23921) in FluoroBrite DMEM. The *E. coli* cells were stained with 1 µM probe for 15 min at 37 °C and imaged without additional washing steps. Imaging was performed with the following laser setup: ex. 488 nm, em. 600/52 nm or em. 542/27 nm, 1.9 mW, 400 ms. Scale bar, 10 µm.

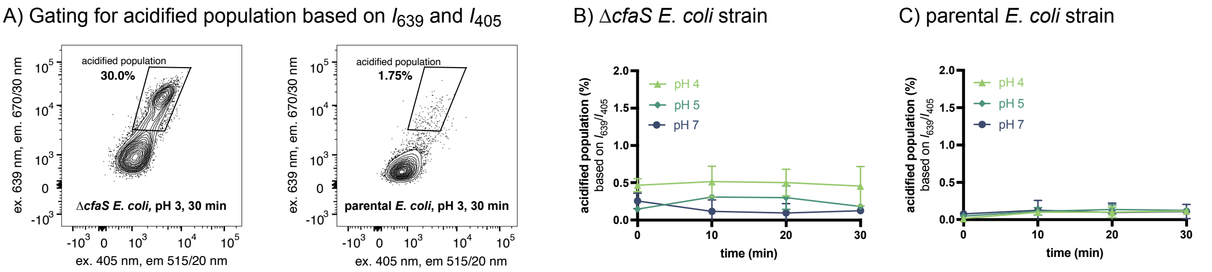

**Figure S14:** Flow Cytometry analysis of parental *E. coli* BW25113 and Δ*cfaS* knockout strain during acid shock in LB. A) Acidified population was gated based on emission intensity of the red ($I_{639}$ > 3.5 × $10^3$) and blue ($I_{405}$ nm > 1 × $10^3$) channels. The bacterial population is represented as a contour plot (5% event density) with outliers shown as dots. B,C) Percentage of acidified population over time under mildly acidic conditions (pH 4, 5, and 7) for Δ*cfaS* (B) and parental *E. coli* strain (C). Data points are means with standard deviation from two biological replicates. All samples were stained with CouCyCF$_3$ **1b** (1 µM) and gated for single bacterial cells.

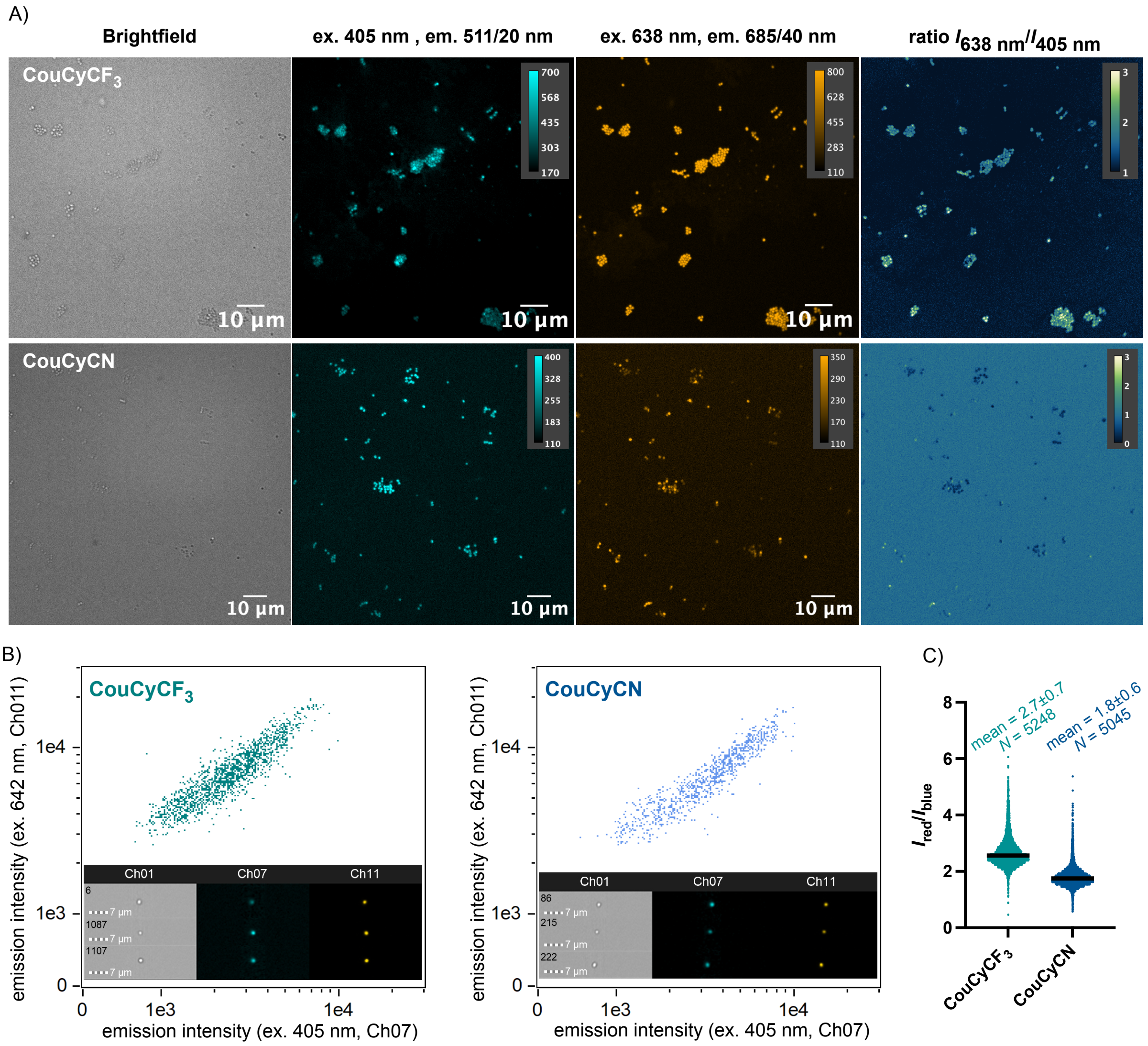

**Figure S15:** Confocal Microscopy and Image Stream experiments with clinical isolate MRSA. A) Microscopy images of MRSA monoculture stained with 2 µM CouCyCF$_3$ **1b** (top panel) or CouCyCN **1c** (lower panel) in 1x PBS (pH 7.4). Imaging was performed with the following laser setup: ex. 405 nm, em. 511/20 nm, 35 mW, 400 ms; ex. 638 nm, em. 685/40 nm, 12 mW, 400 ms. B) Scatter plot and images of MRSA stained with 1 µM CouCyCF$_3$ **1b** (left plot) or CouCyCN **1c** (right plot). C) Scatter plot of MRSA $I_{red}/I_{blue}$ stained with CouCyCF$_3$ **1b** ($N$ = 5248) or CouCyCN **1c** ($N$ = 5045) acquired with Image Stream. Data are summarized as mean with standard deviation. Image Stream was performed with the following setup: Ch01 (Brightfield), Ch07 (ex. 405 nm, em. 435-505 nm, 120 mW), and Ch11 (ex. 642 nm, em. 642-745 nm, 20 mW). Bacterial cells were gated for focused, stained single cells.

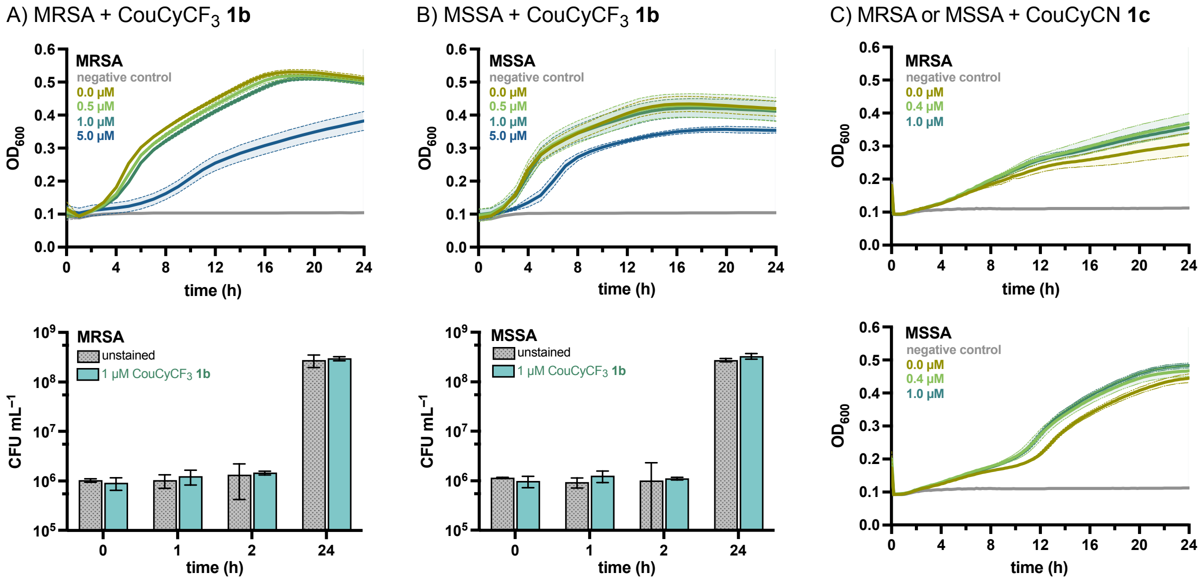

**Figure S16:** Toxicity assay with MRSA and MSSA. A,B) Bacterial growth curve (top) of MRSA (A) and MSSA (B) incubated with $CouCyCF_3$ **1b** (0.5, 1.0, and 5.0 µM). Data points are means of quadruplicates, with the shaded area and dotted line representing standard deviation. Histogram of colony-forming units (CFU) over time (0, 1, 2, and 24 h) at the working concentration of 1 µM $CouCyCF_3$ **1b** (bottom). Bars represent means with standard deviation of two biological replicates. C) Bacterial growth curves of MRSA (top) and MSSA (bottom) incubated with CouCyCN **1c** (0.4 and 1.0 µM). Data points are means of two biological replicates (each quadruplicate), with the shaded area and dotted line representing standard deviation. Bacteria were grown in RPMI + 10% human serum (pH 7.5-7.8). Bacterial cultures were inoculated at $2 \times 10^6$ cells $mL^{-1}$. Medium only served as negative control. CFU was calculated from the number of colonies counted on agar plates.

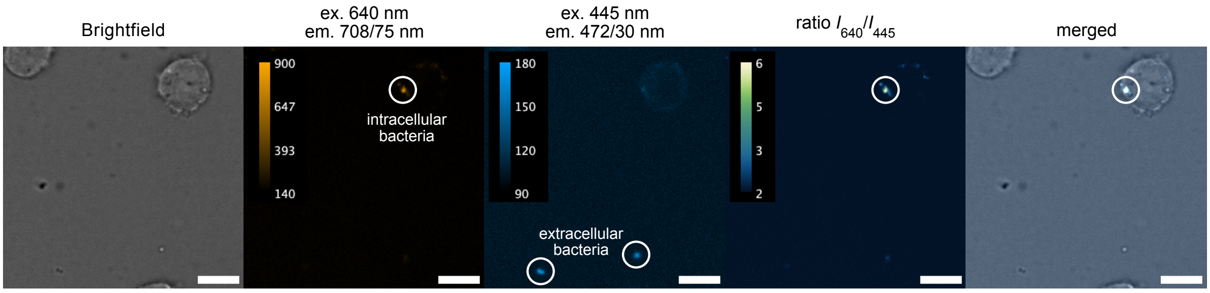

**Figure S17:** Representative confocal microscopy images of *S. epidermidis*-THP-1 coculture. THP-1 monocytes were infected with pre-stained *S. epidermidis* (1 µM $CouCyCF_3$ **1b**) at an MOI of 3. Imaging was performed with the following laser setup: ex. 445 nm, em. 472/30 nm, 1.8 mW, 400 ms; ex. 640 nm, em. 708/75 nm, 3.7 mW, 400 ms. Scale bar, 10 µm.

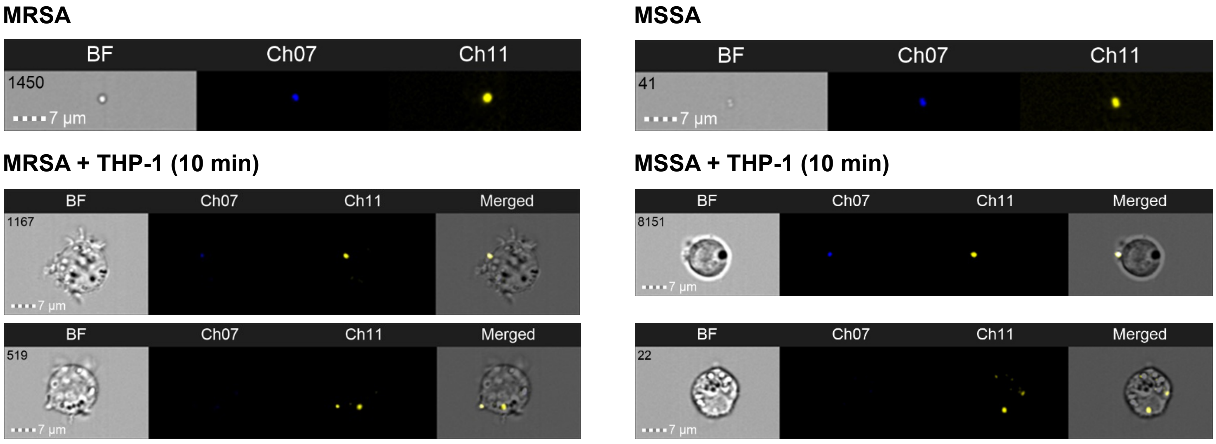

**Figure S18:** Representative images of Image Stream experiment with the clinical isolates MRSA and MSSA. Bacteria were pre-stained with CouCyCF$_3$ **1b** (1 µM) for 10 min, then washed to remove residual dye. THP-1 monocytes were infected, and analysis was performed after 10 min. The following channels were used: BF (Brightfield), Ch07 (ex. 405 nm, em. 435-505 nm), and Ch11 (ex. 642 nm, em. 642-745 nm).

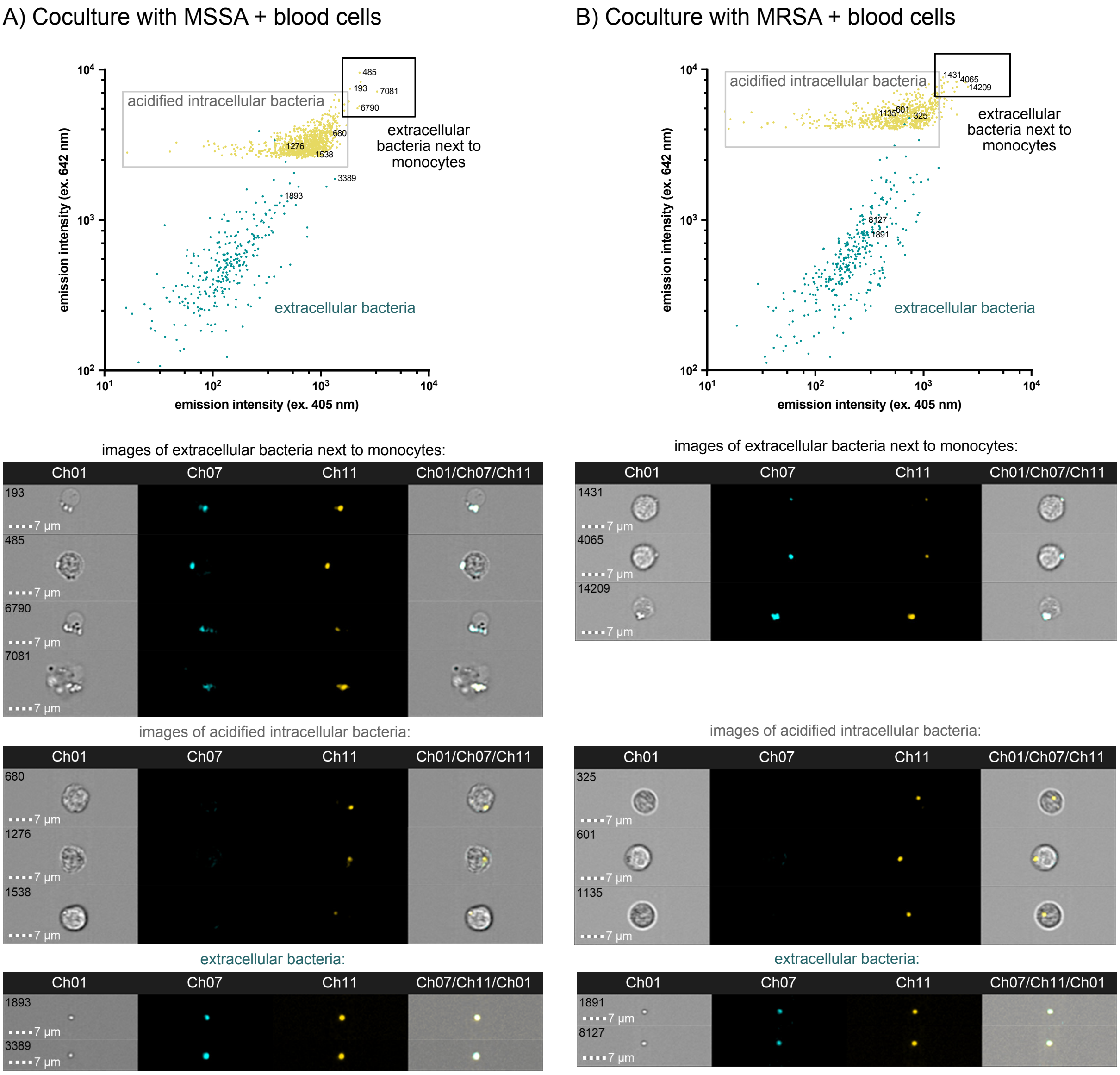

**Figure S19:** Identification of extracellular and intracellular bacteria based on $I_{red}/I_{blue}$ stained with CouCy$CF_3$ **1b** (1 µM). A) Merged scatter plot of MSSA blood cells coculture (yellow) and separately acquired extracellular MSSA data (green; events 1893 and 3389). The main yellow population corresponds to intracellular, acidified bacteria (events 680, 1276, and 1538). The outliers at higher intensities correspond to extracellular bacteria close to blood cells (events 193, 485, 6790, and 7081). B) Scatter plot and images of MRSA blood cells coculture (yellow) and separately acquired extracellular MRSA (green; events 1891 and 8127). The main population corresponds to intracellular, acidified bacteria (events 325, 601, and 1135). The outliers at higher intensities correspond to extracellular bacteria close to blood cells (events 1431, 4065, and 14209). All bacteria were pre-stained, washed, and incubated with blood cells for 10 min at 37 °C before the imaging. Image Stream was performed with the following setup: Ch01 (Brightfield), Ch07 (ex. 405 nm, em. 435-505 nm, 120 mW), and Ch11 (ex. 642 nm, em. 642-745 nm, 120 mW). Data from yellow and green dots were acquired separately and are presented as merged plots. Data were gated for focused, stained, single cells. Internalized bacteria were gated based on the intensity area of Ch11.

#### Supplementary tables

**Table S1:** Calculated p$K_a$ values of $CouCyCF_3$ **1b** and CouCyCN **1c** using the Henderson-Hasselbalch equation $\mathrm{p}K_a = \mathrm{pH} - \left(\log\left(\frac{I_{\max}-I}{I-I_{\min}}\right)\right.$. As intensity $I$, the intensities at the absorbance maxima $A$ (**1b**: $A_{584}$ and $A_{415}$; **1c**: $A_{600}$ and $A_{415}$) or emission maxima $I$ (**1b**: $I_{665}$ and $I_{485}$; **1c**: $I_{675}$ and $I_{505}$) were used.

| | **p$K_a$ (absorbance)** | **p$K_a$ (emission)** |
|---|---|---|
| $CouCyCF_3$ **1b** | 7.0 | 5.7 |
| CouCyCN **1c** | 6.8 | 6.6 |

**Table S2:** Physiochemical properties likely favoring accumulation in Gram-negative bacteria according to the eNTRy guidelines (ionizable Nitrogen, Three-dimensionality, and Rotatable bonds). Properties of $CouCyCF_3$ **1b** and CouCyCN **1c** were calculated with entryway.[23,24]

| | Favored accumulation | $CouCyCF_3$ **1b** | CouCyCN **1c** |
|---|---|---|---|
| **Molecular weight (Da)** | <600-900 Da | 469.5 | 426.5 |
| **Rotatable bonds** | ≤ 5 | 6 | 5 |
| **Globularity** | ≤ 0.250 | 0.04 | 0.03 |
| **Plane of best fit (PBF)** | < 1 | 0.8 | 0.7 |
| **State of charge** | zwitterionic, positive | positive | positive |

**Table S3:** Photophysical properties of $CouCyCF_3$ **1b** (pH 5.5: at 585 nm; pH 8.5: 415 nm; pH 7.4: at 570 nm[13]) and CouCyCN **1c** (pH 5.5: at 600 nm; pH 8.5: at 415 nm; pH 7.4: at 600 nm) in aqueous medium. All values listed are means calculated from preparative triplicates.

| | $\phi_{em}$ in 1x PBS (pH 7.4) | $\phi_{em}$ in 10x PBS (pH 5.5) | $\phi_{em}$ at 415 nm in 10x PBS (pH 8.5) | $\varepsilon$ ($M^{-1}cm^{-1}$) in 1x PBS (pH 7.4) | brightness = $\varepsilon$ x $\phi_{em}$ ($M^{-1}cm^{-1}$) at pH 7.4 |
|---|---|---|---|---|---|
| $CouCyCF_3$ **1b** | 0.8%[13] | 2.0% | <1.0% | 63 640 | 509 |
| CouCyCN **1c** | 1.4% | 1.7% | 4.2% | 51 090 | 715 |

#### References

(1) Newton, G. L.; Arnold, K.; Price, M. S.; Sherrill, C.; Delcardayre, S. B.; Aharonowitz, Y.; Cohen, G.; Davies, J.; Fahey, R. C.; Davis, C. Distribution of Thiols in Microorganisms: Mycothiol Is a Major Thiol in Most Actinomycetes. *J. Bacteriol.* **1996**, *178* (7), 1990–1995. https://doi.org/10.1128/jb.178.7.1990-1995.1996.

(2) Imlay, J. A. The Molecular Mechanisms and Physiological Consequences of Oxidative Stress: Lessons from a Model Bacterium. *Nat. Rev. Microbiol.* **2013**, *11*, 443–454. https://doi.org/10.1038/nrmicro3032.

(3) Tempest, D. W.; Meers, J. L.; Brown, C. M. Influence of Environment on the Content and Composition of Microbial Free Amino Acid Pools. *Microbiol.* **1970**, *64* (2), 171–185. https://doi.org/10.1099/00221287-64-2-171.

(4) Graham, A. I.; Hunt, S.; Stokes, S. L.; Bramall, N.; Bunch, J.; Cox, A. G.; McLeod, C. W.; Poole, R. K. Severe Zinc Depletion of *Escherichia Coli*. *J. Biol. Chem.* **2009**, *284* (27), 18377–18389. https://doi.org/10.1074/jbc.M109.001503.

(5) Groisman, E. A.; Hollands, K.; Kriner, M. A.; Lee, E.-J.; Park, S.-Y.; Pontes, M. H. Bacterial Mg2+ Homeostasis, Transport, and Virulence. *Annu. Rev. Genet.* **2013**, *47*, 625. https://doi.org/10.1146/annurev-genet-051313-051025.

(6) Outten, C. E.; O'Halloran, and T. V. Femtomolar Sensitivity of Metalloregulatory Proteins Controlling Zinc Homeostasis. *Science* **2001**, *292* (5526), 2488–2492. https://doi.org/10.1126/science.1060331.

(7) Ryall, B.; Davies, J. C.; Wilson, R.; Shoemark, A.; Williams, H. D. Pseudomonas Aeruginosa, Cyanide Accumulation and Lung Function in CF and Non-CF Bronchiectasis Patients. *Eur. Respir. J.* **2008**, *32* (3), 740–747. https://doi.org/10.1183/09031936.00159607.

(8) Critchfield, F. E.; Gibson, J. A. Jr.; Hall, J. L. Dielectric Constant for the Dioxane—Water System from 20 to 35°. *J. Am. Chem. Soc.* **1953**, *75* (8), 1991–1992. https://doi.org/10.1021/ja01104a506.

(9) Datsenko, K. A.; Wanner, B. L. One-Step Inactivation of Chromosomal Genes in Escherichia Coli K-12 Using PCR Products. *Proc. Natl. Acad. Sci. U.S.A* **2000**, *97* (12), 6640–6645. https://doi.org/10.1073/pnas.120163297.

(10) Baba, T.; Ara, T.; Hasegawa, M.; Takai, Y.; Okumura, Y.; Baba, M.; Datsenko, K. A.; Tomita, M.; Wanner, B. L.; Mori, H. Construction of Escherichia Coli K-12 in-

Frame, Single-Gene Knockout Mutants: The Keio Collection. *Mol. Syst. Biol.* **2006**, *2*, 2006.0008. https://doi.org/10.1038/msb4100050.

(11) Zarkan, A.; Caño-Muñiz, S.; Zhu, J.; Al Nahas, K.; Cama, J.; Keyser, U. F.; Summers, D. K. Indole Pulse Signalling Regulates the Cytoplasmic pH of E. Coli in a Memory-Like Manner. *Sci. Rep.* **2019**, *9* (1), 3868. https://doi.org/10.1038/s41598-019-40560-3.

(12) An, K. L.; Shin, S. R.; Jun, K.; Park, S. Y. The Synthesis and Light Absorption Behaviour of Novel Coumarin Chromophores. *J. Korean Chem. Soc.* **2014**, *58* (3), 297–302. https://doi.org/10.5012/jkcs.2014.58.3.297.

(13) Tirla, A.; Rivera-Fuentes, P. Development of a Photoactivatable Phosphine Probe for Induction of Intracellular Reductive Stress with Single-Cell Precision. *Angew. Chem. Int. Ed.* **2016**, *55* (47), 14709–14712. https://doi.org/10.1002/anie.201608779.

(14) Wolf, N.; Kersting, L.; Herok, C.; Mihm, C.; Seibel, J. High-Yielding Water-Soluble Asymmetric Cyanine Dyes for Labeling Applications. *J. Org. Chem.* **2020**, *85* (15), 9751–9760. https://doi.org/10.1021/acs.joc.0c01084.

(15) Owens, E. A.; Bruschi, N.; Tawney, J. G.; Henary, M. A Microwave-Assisted and Environmentally Benign Approach to the Synthesis of near-Infrared Fluorescent Pentamethine Cyanine Dyes. *Dyes Pigm.* **2015**, *113*, 27–37. https://doi.org/10.1016/j.dyepig.2014.07.035.

(16) Martin, A.; Rivera-Fuentes, P. A General Strategy to Develop Fluorogenic Polymethine Dyes for Bioimaging. *Nat. Chem.* **2024**, *16* (1), 28–35. https://doi.org/10.1038/s41557-023-01367-y.

(17) Kim, I. K.; Erickson, K. L. Models for Uleine-Alkaloid Biogenesis. *Tetrahedron* **1971**, *27* (17), 3979–3991. https://doi.org/10.1016/S0040-4020(01)98123-2.

(18) Liang, Z.; Sun, Y.; Duan, R.; Yang, R.; Qu, L.; Zhang, K.; Li, Z. Low Polarity-Triggered Basic Hydrolysis of Coumarin as an AND Logic Gate for Broad-Spectrum Cancer Diagnosis. *Anal. Chem.* **2021**, *93* (36), 12434–12440. https://doi.org/10.1021/acs.analchem.1c02591.

(19) Checa, M.; Millan-Solsona, R.; Blanco, N.; Torrents, E.; Fabregas, R.; Gomila, G. Mapping the Dielectric Constant of a Single Bacterial Cell at the Nanoscale with Scanning Dielectric Force Volume Microscopy. *Nanoscale* **2019**, *11* (43), 20809–20819. https://doi.org/10.1039/C9NR07659J.

(20) Yoon, S. A.; Cha, S. H.; Jun, S. W.; Park, S. J.; Park, J.-Y.; Lee, S.; Kim, H. S.; Ahn, Y. H. Identifying Different Types of Microorganisms with Terahertz Spectroscopy. *Biomed. Opt. Express* **2019**, *11* (1), 406–416. https://doi.org/10.1364/BOE.376584.

(21) Esteban-Ferrer, D.; Edwards, M. A.; Fumagalli, L.; Juárez, A.; Gomila, G. Electric Polarization Properties of Single Bacteria Measured with Electrostatic Force Microscopy. *ACS Nano* **2014**, *8* (10), 9843–9849. https://doi.org/10.1021/nn5041476.

(22) invitrogen. Molecular Probes$^{TM}$ Handbook A Guide to Fluorescent Probes and Labeling Technologies - Chapter 20: pH Indicators. In *Molecular Probes$^{TM}$ Handbook A Guide to Fluorescent Probes and Labeling Technologies - Chapter 20: pH Indicators*; 2010; pp 883–902.

(23) Richter, M. F.; Drown, B. S.; Riley, A. P.; Garcia, A.; Shirai, T.; Svec, R. L.; Hergenrother, P. J. Predictive Compound Accumulation Rules Yield a Broad-Spectrum Antibiotic. *Nature* **2017**, *545* (7654), 299–304. https://doi.org/10.1038/nature22308.

(24) Ropponen, H.-K.; Richter, R.; Hirsch, A. K. H.; Lehr, C.-M. Mastering the Gram-Negative Bacterial Barrier – Chemical Approaches to Increase Bacterial Bioavailability of Antibiotics. *Adv. Drug Deliv. Rev.* **2021**, *172*, 339–360. https://doi.org/10.1016/j.addr.2021.02.014.

### Appendix

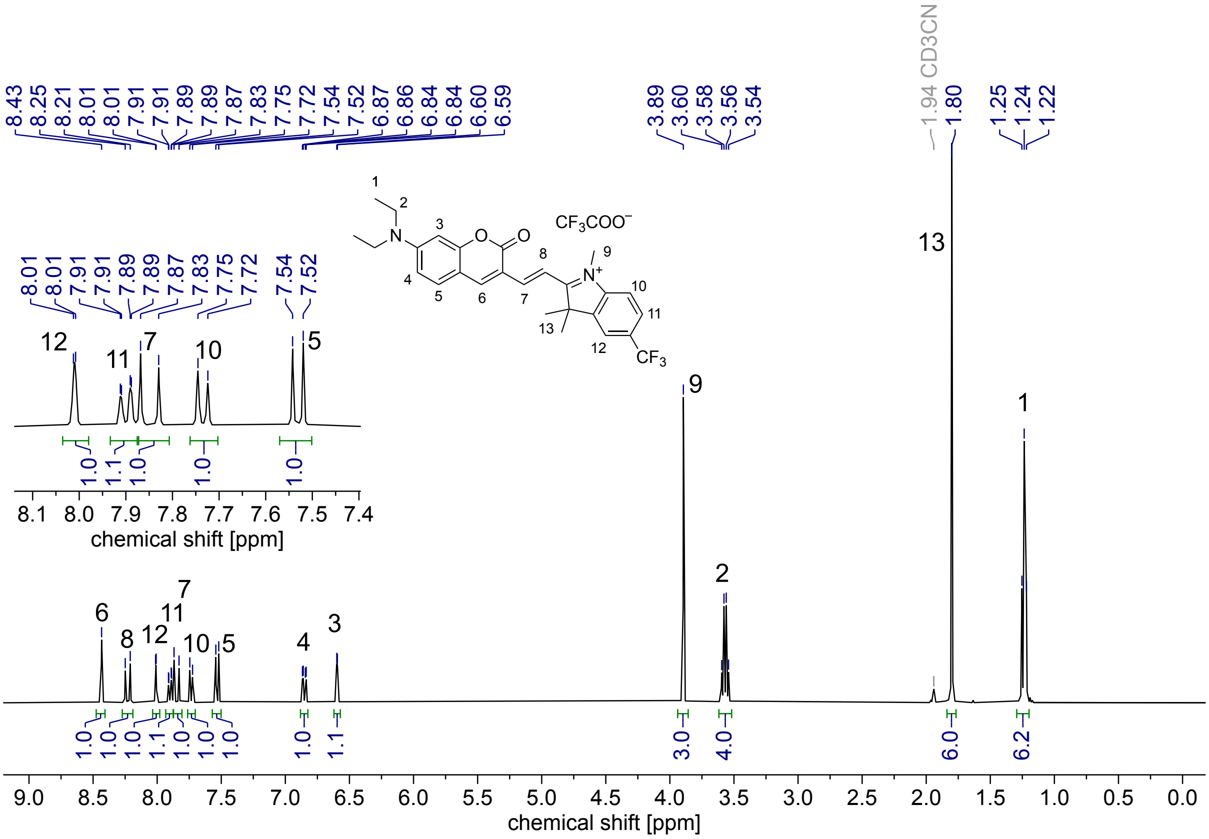

**Figure S20:** $^1$H-NMR (400 MHz, $CD_3CN$) of CouCyCF$_3$ **1b**.

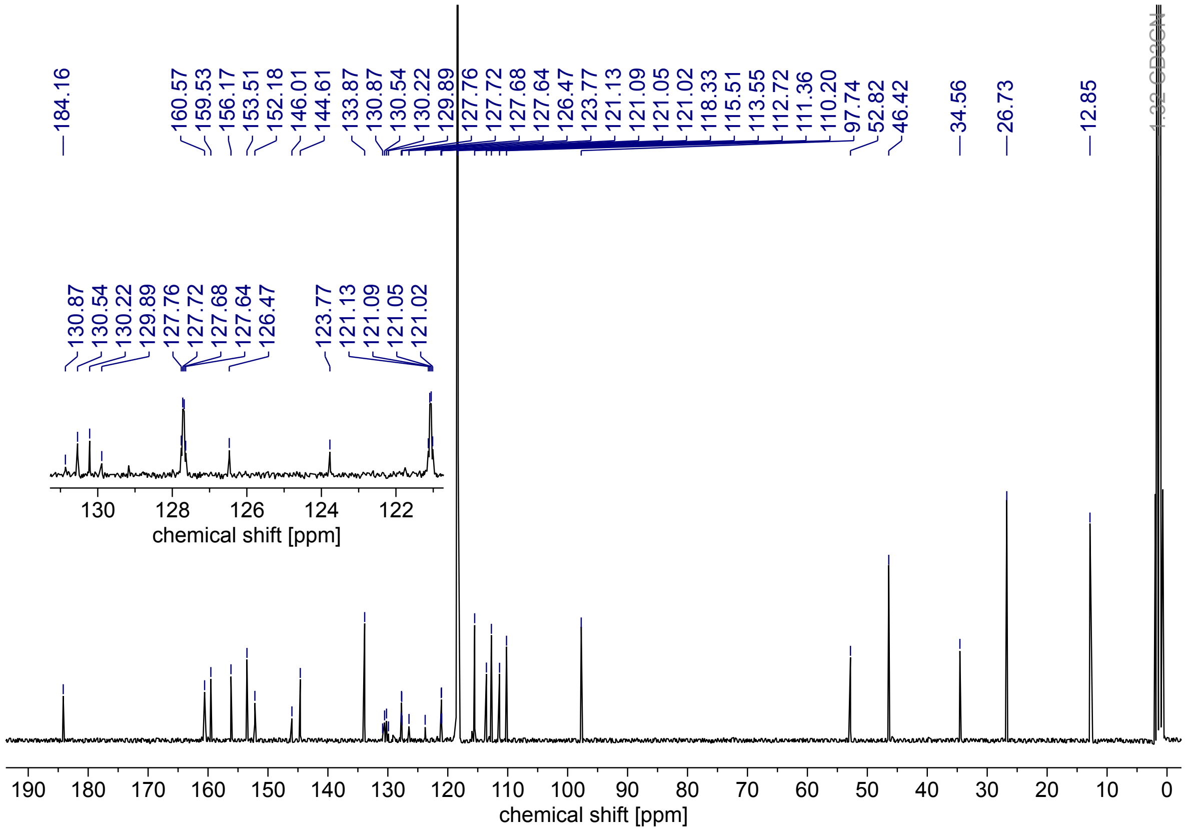

**Figure S21:** $^{13}$C-NMR (101 MHz, $CD_3CN$) of CouCyCF$_3$ **1b**.

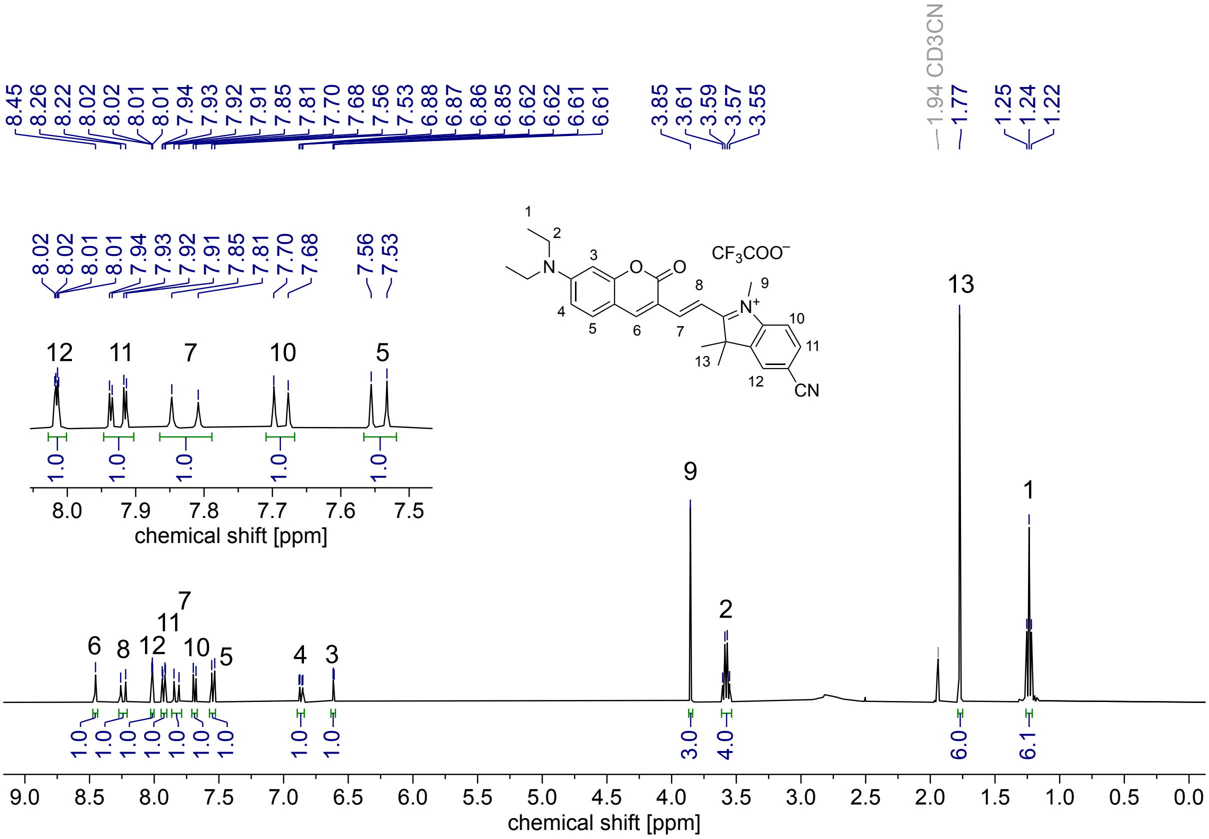

**Figure S22:** $^1$H-NMR (400 MHz, $CD_3CN$) of CouCyCN **1c**.

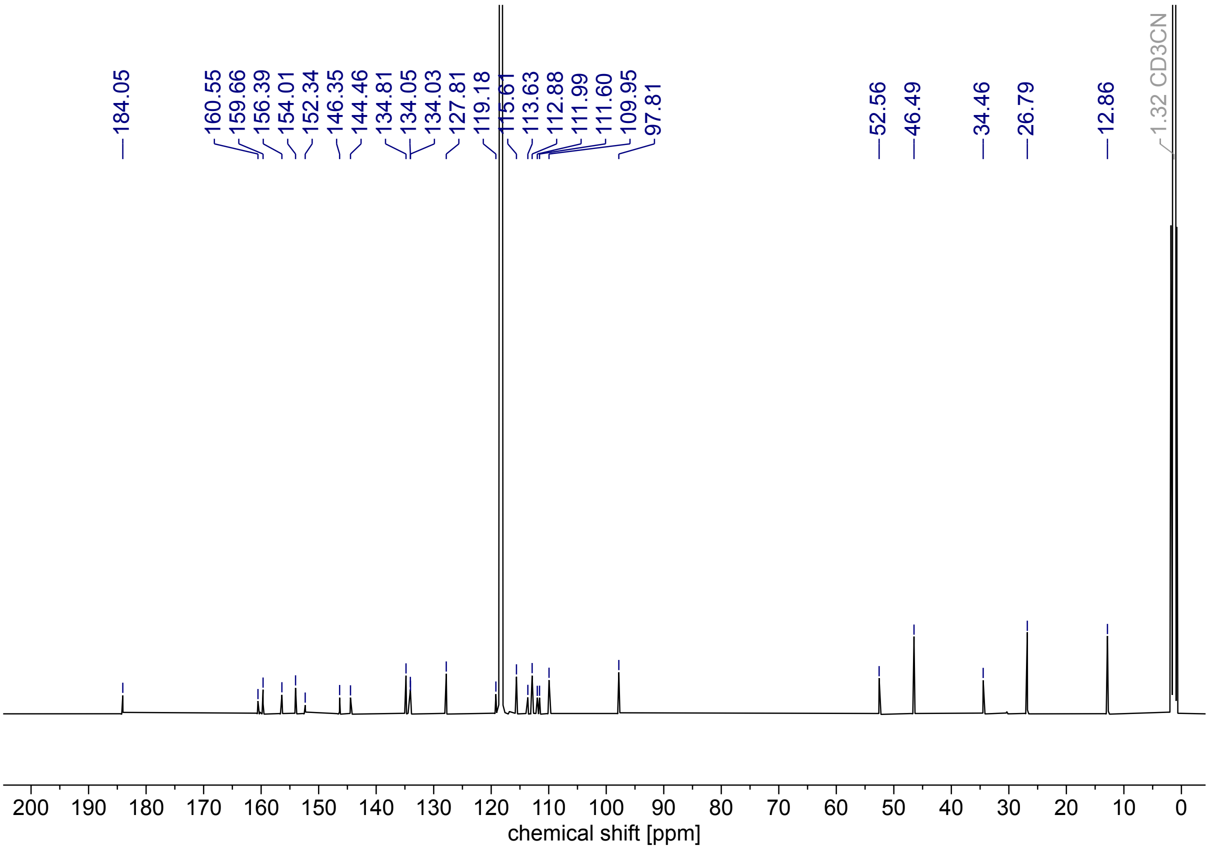

**Figure S23:** $^{13}$C-NMR (126 MHz, $CD_3CN$) of CouCyCN **1c**.
